# Mammalian condensin I controls higher-order chromosome organization and homologous recombination in meiotic prophase I

**DOI:** 10.64898/2026.09.15.751738

**Authors:** Laurine Dal Toe, Boubou Diagouraga, Alexandre Nore, Audrey Bost, Estelle Grosjean, Julien Cau, Thomas Robert

## Abstract

The meiotic chromosome structure, defined by the axis-chromatin loop organization, is essential for proper homologue synapsis through the formation of the synaptonemal complex, and faithful homologous recombination. Here, we show that the conserved SMC-condensin I complex, which shapes the genome through chromatin loop extrusion, plays a pivotal role in controlling the higher- order prophase I meiotic chromosome structure. By analyzing the NCAPD2 condensin I subunit, we found that this complex colocalizes with prophase I chromatin and is essential for male fertility. NCAPD2 restricts chromatin loop extension and controls the localization of chromosome structural proteins, including the axial component HORMAD1 and central element proteins (SYCP1, TEX12, SIX6OS1), thus regulating the synaptonemal complex width. Moreover, NCAPD2 promotes timely and efficient recombination by controlling γH2AX, DMC1 and pro-crossover proteins turnover. We propose that condensin I, by organizing chromatin loops in prophase I, controls synaptonemal complex organization and recombination outcome in mammals.

## Introduction

During meiosis, the proper segregation of the homologous chromosomes relies on two pivotal interdependent molecular events that promote their physical connection: i) the formation of a meiosis-specific higher-order chromosome organization and ii) DNA homologous recombination (HR) ^1^.

Meiotic recombination initiates at prophase I onset with the programmed formation of DNA double- strand breaks (DSBs) at specific genomic locations, the hotspots ^2–4^. Following their subsequent processing, a fraction of these breaks will be converted into double Holliday Junctions (dHJ) and resolved as crossovers (CO), which correspond to the future physical connection points between homologues (i.e. chiasmata) ^1,5^. Meiotic HR occurs in the context of a complete structural reorganization of the chromosomes, highlighted by the formation of the synaptonemal complex (SC), a protein structure that connects the homologues along their entire length ^6^. SC formation starts in leptonema, by the progressive assembly of the axial elements, on which both sister chromatids are anchored, forming chromatin loops. DSBs form at this step. A common view is that DSB prone- sequences lie in the chromatin loops, suggesting that their organization (*e.g.* size, distribution) governs DSB patterning^6–10^. At zygonema, synapsis starts through the attachment of the transverse filaments to each axis, now named lateral elements, and the connection of the central element to the transverse filaments, which progressively connect and align the two homologues. At pachynema, the SC is fully assembled and the two homologues are fully synapsed ^11^. SC organization also plays a critical role in CO regulation. Pro-CO proteins are localized within the central part of the SC indicating the presence of DNA recombination intermediates at this location ^1,5,11^. Pro-CO proteins stabilizes dHJs, and different models propose that SC organization influences their distribution/numbers, and thus CO outcome ^5,12–15^.

At the protein level, the synapsis of homologs is dependent upon the prior formation of the axis that is formed by the interaction of cohesin complexes (composed of the meiotic specific REC8/RAD21L kleisin, SMC1β and STAG3 subunits and ubiquitous subunits) and meiosis-specific proteins, such as SYCP3/SYCP2 fibrillar complex and the HORMA domain proteins HORMAD1/2 ^10,11^. HORMAD1 is progressively depleted from the axis upon SC central element formation and acts as a regulatory unit between the SC structure and HR, by controlling DSB formation and monitoring proper synapsis and HR ^16–19^. The transverse filament is formed through SYCP1 self-assembly as a head-to-head tetramer that connects the lateral elements to the central element, particularly to the SYCE3 protein that stabilizes SYCP1 assembly ^11^. Four additional proteins compose the central element in mammals: SYCE1, SIX6OS1, SYCE2 and TEX12 ^11^. Although the mammalian SC composition starts to be better characterized, how the meiotic chromatin architecture is maintained throughout prophase I, in the context of the SC, and whether it controls HR remains unclear.

Among the proteins known to control the higher-order chromosome organization, SMC-condensin complexes are conserved multi-subunit complexes, described as the universal organizers of chromosomes ^20^. In mammals, two condensin complexes have been described: condensin I and II. Both are composed of two Structural Maintenance of Chromosome (SMC) proteins (SMC2 and SMC4) that assemble in a V-shaped structure. The vertices of this structure are connected by a kleisin subunit (NCAPH and NCAPH2 for condensin I and II, respectively), resulting in a closed ring structure that can encircle DNA. Kleisin subunits recruit two additional HEAT repeat regulatory subunits (NCAPD2/G and NCAPD3/G2 for mammalian condensin I and II, respectively) ^21^. Condensin I and II share redundant functions, but also have specific roles, suggesting that both are essential ^20,22^. Condensins act primarily in metaphase and anaphase of mitotic and meiotic cell divisions, where they promote chromosome compaction and individualization (condensation), which are essential for segregation ^22,23^. More specifically, it was proposed that condensin I promotes lateral chromosome compaction, while condensin II promotes longitudinal chromosome compaction ^21,24^. Biophysical *in vitro* experiments demonstrated that condensins form DNA loops through the loop extrusion process ^21,25^.

In meiotic prophase I, condensin function is less understood. In yeast, it constrains the axis length and promotes the proper recruitment of the Hop1, Red1 axial proteins and Zip1 central element protein (respectively HORMAD1, SYCP2 and SYCP1 mouse orthologues), as well as cohesin removal at the end of prophase I ^15,26–28^. DSB formation is condensin-independent, but condensin is required to resolve recombination-dependent chromosome linkages at prophase I end, and to suppress recombination at the rDNA locus ^26,27,29^. More recently, it was shown that condensins influence CO number, and are enriched at CO designation sites, promoting the local reorganization of the axis composition ^15,28,30^. In nematodes, the condensin I subunit DPY-28 (NCAPD2 orthologue) regulates DSB formation and CO distribution, potentially by controlling axis length, through a functional interaction with the HIM-3 HORMAD1 orthologue ^31–34^. Although condensin function in SC protein assembly was shown in yeast and nematodes, their implication in meiotic chromatin loop organization remains unknown, as well as their role in these processes in mammalian prophase I.

Here, we performed an extensive characterization of the condensin I complex role, particularly the NCAPD2 subunit, in mouse prophase I. We found that this protein localizes predominantly on chromatin throughout nucleus rather than on axis chromosome structure, and is required for proper spermatogenesis and fertility, by controlling meiotic higher-order chromosome organization. Condensin I promotes the compaction of prophase I chromatin, potentially by limiting chromatin loop extension. NCAPD2 also controls the SC superstructure, limits its width by controlling the HORMAD1 pattern and central element protein recruitment, but without affecting axis length. Lastly, NCAPD2 promotes proper HR, by favouring the recombination protein turnover and final resolution of recombination intermediates. We propose that in mammalian meiotic prophase I, condensin I regulates the higher-order chromosome organization and HR outcome.

## Results

### The NCAPD2 condensin I subunit colocalizes with chromatin, throughout prophase I

To better understand the expression of the condensin I complex during meiotic prophase I in the mouse, we analysed NCAPD2 spatiotemporal pattern of recruitment in prophase I spermatocytes of wild type (WT) C57BL/6J mice. First, we confirmed NCAPD2 expression in early prophase I nuclei by western blotting of whole protein extracts from WT juveniles’ testis cells with synchronized spermatogenesis to obtain samples enriched in leptotene/zygotene spermatocytes (see ^8^ for meiotic synchronization procedure) (Figure 1A). Remarkably, the detection of the SMC2 and NCAPD3 subunits also indicates the presence of both condensin I and II complexes at these early prophase I stages. Second, we performed a time course analysis of NCAPD2 expression by western blotting using testis nuclear protein extracts from juveniles (from 3 dpp to 21 dpp) and adult WT mice (60 dpp and 148 dpp). NCAPD2 expression peaked at 6-12 dpp, corresponding to meiotic prophase I onset, and was strongly reduced in extracts from adult mice (Figure 1B). Lastly, immunofluorescence analysis of NCAPD2 nuclear localization on chromosome spreads from 18 dpp juvenile WT spermatocytes showed many NCAPD2 foci that colocalized with DAPI-stained chromatin, from leptonema to diplonema (Figure 1C and Figure S1A). These findings suggest that, unlike SMC- cohesin complexes which specifically localize on chromosome axis ^10^, NCADP2 decorates meiotic spermatocyte chromatin throughout the nucleus in prophase I.

**Figure 1.**
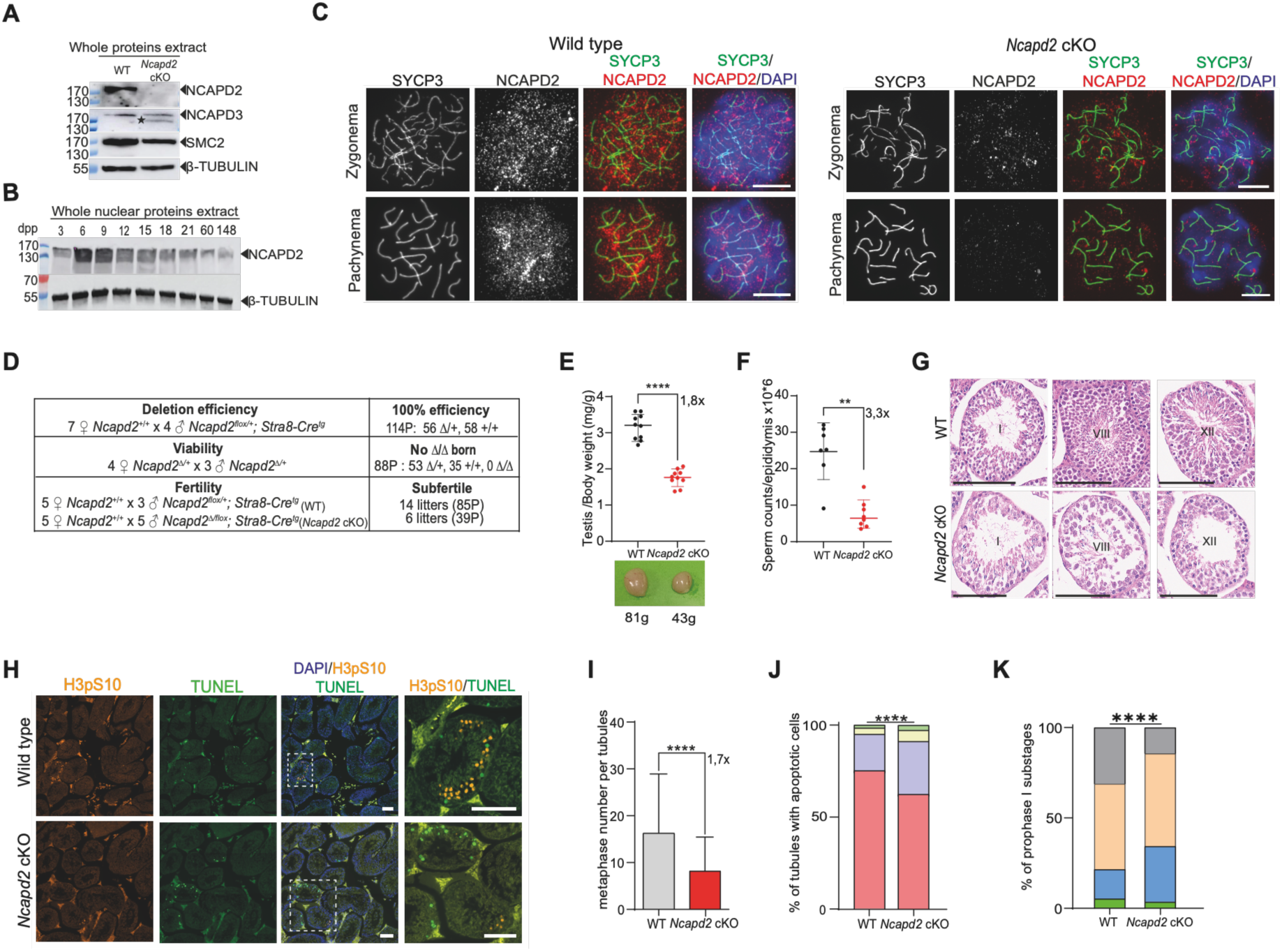
NCAPD2 colocalizes with chromatin in prophase I and is required for meiotic progression. (A) Immunodetection of NCAPD2, NCAPD3 and SMC2 in meiocytes by western blotting of whole protein extracts from testes of wild type (WT) and *Ncapd2* cKO synchronized mice; β-tubulin was used as loading control (n=3 for NCAPD2 and SMC2 and n=2 for NCAPD3); asterisk, non-specific band. (B) Immunodetection of NCAPD2 during the first wave of meiosis by western blotting of nuclear protein extracts from testes of WT mice (3 to 148 dpp); β-tubulin was used as loading control. (C) NCAPD2 immunolocalization in meiocytes. Representative images of SYCP3 (green), NCAPD2 (red) and DAPI (blue) immunostaining in zygotene and pachytene nuclei of WT and *Ncapd2* cKO mice at 18 dpp (scale bar, 10 µm). (D) Crosses to determine the deletion efficiency of the flox allele (4 independent crosses of one male with one or two females), the viability of homozygous Δ mice (3 independent crosses of one male with one or two females), and fertility of *Ncapd2* cKO males (3 crosses, one male with one or two females for the wild type, and 5 crosses, one male with one female for *Ncapd2* cKO). Total number of animals is given in the table. (E) Testis weight is reduced in *Ncapd2* cKO mice. *(Top)* Quantification of the mean testis weight relative to the body weight (mg/g) in WT and *Ncapd2* cKO mice at 60 dpp. Each dot represents one testis (n=10). *(Bottom)* Representative images of WT and *Ncapd2* cKO testes at 60 dpp. (F) Sperm production is reduced in *Ncapd2* cKO mice. Sperm count per epididymis; each dot represents the mean sperm number from both testes of one mouse at 60 dpp (n=7 for WT and 8 for *Ncapd2* cKO). (G) Haematoxylin and eosin-stained sections of testes from 60 dpp WT and *Ncapd2* cKO mice (scale bar, 100 µm). Representative images at stages I, VIII, and XII are shown. (H) Detection of apoptosis and metaphases in testis sections. Representative images of testis sections from 60 dpp WT and *Ncapd2* cKO mice labelled with TUNEL (green), H3pS10 (orange), and DAPI (blue). White boxes indicate regions shown at higher magnification (scale bar, 250 µm; 100 µm in zoom). (I) Number of metaphase cells per stage XII tubule in testis sections from 60 dpp WT and *Ncapd2* cKO mice (number of stages XII tubules: 74 for WT and 83 for *Ncapd2* cKO). (J) Apoptotic cells are increased in *Ncapd2* cKO testes. Graph showing the percentage of seminiferous tubules containing 0 (red), < 3 (blue), between 4 and 6 (yellow), or > 6 (green) apoptotic cells in testis sections from 60 dpp WT and *Ncapd2* cKO mice (mean of 3 independent experiments with 974, 1292, 1127 WT tubules and 684, 1883, 1109 *Ncapd2* cKO tubules). (K) Prophase I progression depends on NCAPD2. Graph showing the percentage (mean of 3 independent experiments) of spermatocyte nuclei at each prophase I substage in 18 dpp WT and *Ncapd2* cKO mice; green, leptonema (L); blue, zygonema (Z); yellow, pachynema (P); and grey, diplonema (D). Number of nuclei: wild type L 7, 6, 4; Z: 15, 23, 14; P: 34, 56, 61 and D: 45, 23, 31; *Ncapd2* cKO L: 2, 3, 6; Z 32, 44, and 19; P: 66, 36, 56 and D: 6, 16, and 22. P values were determined with the two-tailed unpaired Mann-Whitney test in (e), (f) and (h) and the χ2 test in (j) and (k); black or red bars show the mean values ± SD.

### Generation of a mouse strain in which *Ncapd2* is specifically knocked out in prophase I

To investigate NCAPD2 functional role in prophase I, we generated a conditional knockout mouse line, in which *Ncapd2* was specifically ablated at meiosis I onset. We first generated a line containing a floxed allele of the *Ncapd2* gene: *Ncapd2^flox^*(Figure S1B). The LoxP sites flanked exons 7 and 9 and the deletion led to a frameshift in exon 10 (*Ncapd2^Δ^*) that truncates the 1153 C-terminal residues (total protein length = 1392 aa) (Figure S1C, D). As this truncated region contains most of the HEAT repeat motifs ^35^, it should behave as a null allele. Then, to specifically inactivate the gene in the male germline at meiotic prophase I onset, we used a transgene that expresses the Cre recombinase under the control of the *Stra8* promoter ^36^. The Cre-mediated deletion efficiency, estimated by genotyping the progeny of heterozygous *Ncapd2^flox/+^ Stra8-Cre^tg^* males crossed with WT females, was 100% (n=172 pups) (Figure 1D and S1E, F for the genotyping strategy). When we mated the resulting *Ncapd2^Δ/+^* males and females, we never obtained *Ncapd2^Δ/Δ^* mice (Figure 1D), confirming that *Ncapd2* is essential for embryonic development.

We confirmed NCAPD2 depletion in prophase I *Ncapd2^Δ/flox^ Stra8-Cre^tg^* spermatocytes (named *Ncapd2* cKO hereafter) by two independent approaches. First, western blot analysis of whole protein extracts from testes with synchronized spermatogenesis from juvenile mice showed that unlike in WT samples, NCAPD2 was barely detectable in *Ncapd2* cKO leptotene/zygotene cells (Figure 1A). The similar expression level of SMC2 and NCAPD3 in WT and *Ncapd2* cKO spermatocytes suggests that NCAPD2 absence affects specifically the condensin I complex (Figure 1A). Second, immunostaining experiments revealed a significant loss of NCAPD2 signal in 80%-70% of foci at leptonema, zygonema and pachynema in *Ncapd2* cKO nuclei compared with WT nuclei (Figure 1C and S1A, G). At these stages, in 20%-30% of nuclei, NCAPD2 signal was similar to WT, suggesting incomplete or delayed Cre-mediated excision, as previously reported ^37–39^. The percentage of *Ncapd2* cKO nuclei exhibiting a NCAPD2 signal similar to WT increased to ∼70% at diplonema, suggesting that the majority of NCAPD2-depleted nuclei cannot reach late prophase I stages, beyond pachytene. However, the 100% deletion efficiency found in the progeny revealed that the Cre-mediated excision and protein depletion will ultimately occur at diplonema or later during spermatogenesis. Consequently, in the rest of the study, unless otherwise stated, the *Ncapd2* cKO diplotene nuclei, and following stages, will not be discussed (see Supplementary discussion).

### NCAPD2 is required for normal prophase I progression

*Ncapd2* cKO males developed normally and reached adulthood. To determine whether NCAPD2 is required for normal male meiosis completion, we analysed the fertility of *Ncapd2* cKO males by crossing them with wild type females. After six months of breeding, we observed an important reduction of the fertility associated to the mutation (Figure 1D). This phenotype, the 1.8-fold decrease in testis weight and the 3-fold decrease in sperm count in *Ncapd2* cKO compared with WT adult males (Figures 1E, F) showed a defect in spermatogenesis, leading to subfertility in the absence of NCAPD2.

Histological analysis of haematoxylin-eosin-stained testis sections from *Ncapd2* cKO males revealed that the seminiferous tubules (all spermatogenesis stages, from I to XII) (Figure 1G) were completely disorganized, with a drastic reduction of cell number. In line with this observation, H3pS10 immunostaining of testis sections showed a 1.7-fold decrease in the number of metaphase cells per tubule and, by TUNEL assay, we found a 13% increase in the proportion of tubules containing apoptotic cells in *Ncapd2* cKO *vs* WT animals (Figure 1H-J). These results confirmed a spermatogenesis defect in *Ncapd2* cKO males. Moreover, apoptotic cells in *Ncapd2* cKO testes were mainly located internal to the most peripheral cells (namely the spermatogonia), which is compatible with a prophase I defect (higher magnification in Figure 1H). Although *Ncapd2* cKO tubules exhibited significant disruption, we detected spermatozoa with normal morphology in the sperm and testis sections (Figure 1F, G), in agreement with the potential incomplete or delayed Cre-mediated excision (see Supplementary discussion).

We then monitored prophase I progression by co-staining surface-spread spermatocyte nuclei with antibodies against SYCP3 and SYCP1 (SC axial protein and lateral element protein, respectively), to follow SC formation, and against γH2AX (DSB formation marker). In *Ncapd2* cKO spermatocytes, we observed an increase in the proportion of zygotene/pachytene cells (82% *vs* 64% in WT spermatocytes) and decrease in diplotene cells (14% *vs* 31% in WT spermatocytes) (Figure 1K). In *Ncapd2* cKO samples, pachytene cells might represent the majority of NCAPD2-depleted cells, while the small proportion of diplotene cells might correspond to NCAPD2-expressing cells, because of the proposed incomplete activity of the Cre-mediated deletion system (see Supplementary discussion).

Altogether, these results show that NCAPD2 is required for normal prophase I progression, and are in accordance with a role for this condensin I subunit in mouse prophase I.

### NCAPD2 controls chromatin compaction during prophase I

Chromatin occupancy in prophase I is constrained by the organization of chromatin in loops, anchored along a protein axis ^40^ (Figure 2A). As condensin I was proposed to promote the chromosome lateral compaction during mitosis by generating DNA loops ^21^, and because we detected NCAPD2 on chromatin in prophase I, we asked whether condensin I contributes to chromatin organization and loop formation at this stage.

**Figure 2.**
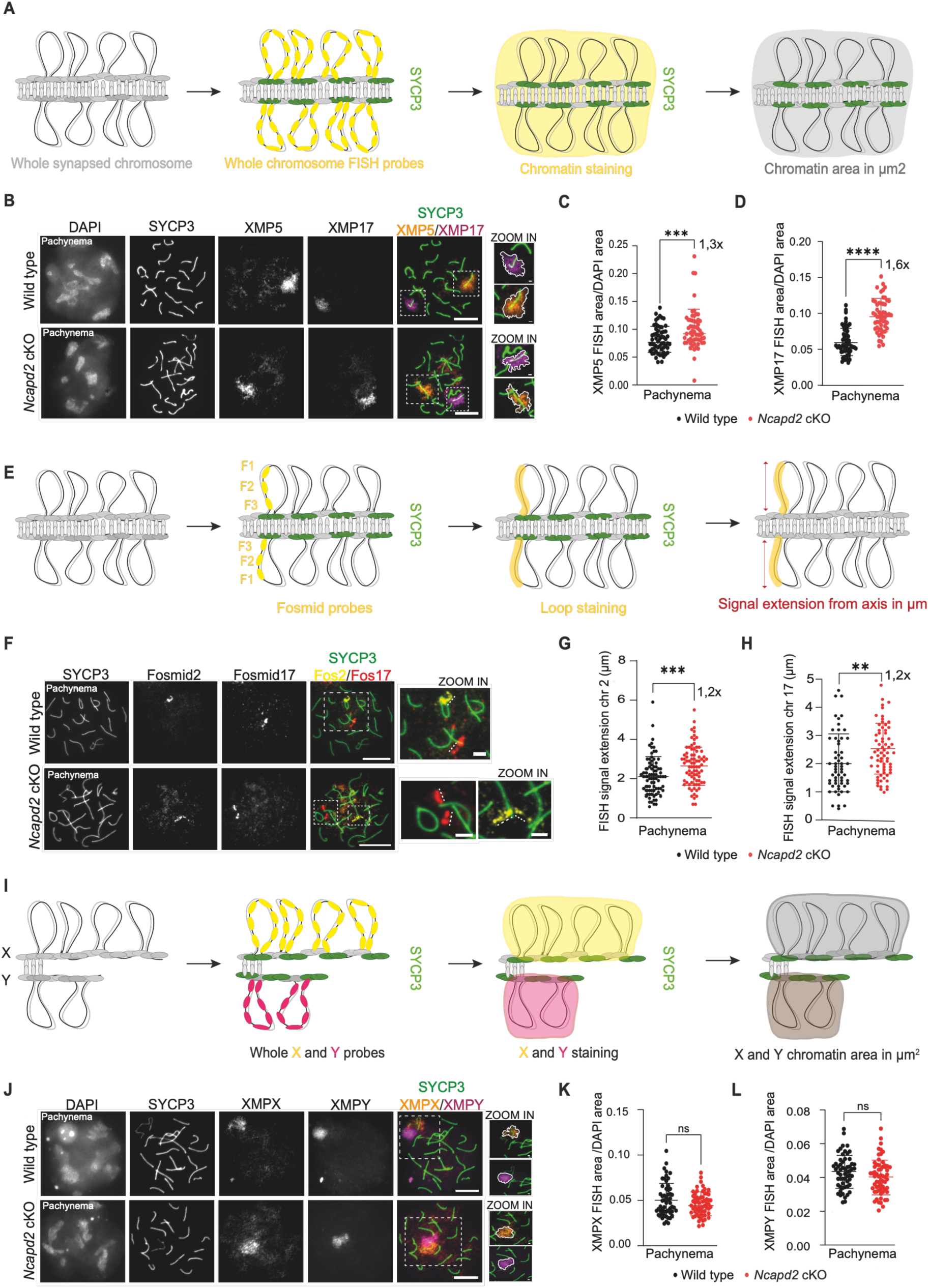
NCAPD2 promotes chromatin compaction on the autosomes 2, 5 and 17. (A) Experimental set-up of the immuno-FISH experiments on autosomes using whole chromosome probes and an anti-SYCP3 antibody to stain the lateral element. (B) Whole autosome chromatin staining. Representative images of WT and *Ncapd2* cKO pachytene nuclei stained with the anti-SYCP3 antibody (green) and labelled with FISH probes for chromosomes 5 (XMP5, orange) and 17 (XMP17, magenta). (C and D) NCAPD2 controls chromatin compaction of the whole chromosomes 5 and 17. Ratio of the XMP5 (C) or XMP17 (D) FISH probe area relative to the DAPI area in WT and *Ncapd2* cKO pachytene nuclei. Number of pachytene nuclei: 60 for both XMP5 and XMP17 (WT); 59 and 55 for XMP5 and XMP17 (*Ncapd2* cKO). (E) Experimental set-up of the immuno-FISH experiment on autosomes using fosmid probes. Three consecutive fosmid FISH probes are hybridized and an anti-SYCP3 antibody is used to measure the FISH signal extension (yellow) from the axis in µm (red arrow). (F) Chromatin signal extension on chromosomes 2 and 17. Representative images of SYCP3 immunostaining (green) and FISH labelling of chromosomes 2 (yellow) and 17 (red) in pachytene nuclei of WT and *Ncapd2* cKO mice. (G and H) NCAPD2 controls chromatin extension from the axis on chromosomes 2 and 17. Quantification of the FISH signal extension (in µm) on chromosomes 2 (G) and 17 (H) in pachytene nuclei from WT and *Ncapd2* cKO. Number of nuclei stained with fosmid 2 (n=81 for both WT and *Ncapd2* cKO), and stained with fosmid 17 (n=58 for WT and 67 for *Ncapd2* cKO). (I) Experimental set up of the immuno-FISH experiment on the sex chromosomes using whole chromosome FISH probes, and SYCP3 immunostaining. The FISH signals surround the chromosomes X (yellow) and Y (magenta). (J) Whole X-Y chromosome chromatin staining. Representative images of WT and *Ncapd2* cKO pachytene nuclei immunostained with an anti-SYCP3 antibody (green) and labelled with the FISH probes for chromosomes X (XMPX, orange) and Y (XMPY, magenta). (K and L) X-Y whole chromosome chromatin compaction is NCAPD2-independent. Ratio of the XMPX (K) and XMPY (L) FISH probe area relative to the DAPI area in pachytene nuclei from WT and *Ncapd2* cKO testes. Numbers of pachytene nuclei for WT and *Ncapd2* cKO: 55 for XMPX and 62 for XMPY. For the experiments shown in this figure, 18 dpp mice were used. White boxes indicate areas at higher magnification where the measured area is marked with white lines (B and J) and signal extension with dashed lines (F). Scale bars, 10 µm and 2 µm for the zoomed images. P values were determined using the two-tailed unpaired Mann– Whitney test. Black or red bars, mean values ± SD.

To assess chromatin occupancy at the scale of the entire bivalent chromosomes, when the SC is fully assembled (pachytene stage), we performed fluorescence in situ hybridization (FISH) using whole chromosome FISH probes for the autosomes 2, 5, 8 and 17 on chromosome spreads from WT and *Ncapd2* cKO pachytene nuclei. The size of these chromosomes ranges from ∼95 Mb to ∼181 Mb, thus covering the spectrum of mouse chromosome sizes. Then, we measured the area in µm^2^ of the FISH signals and normalized them to the DAPI nuclear area in each nucleus, to account for potential spreading differences among nuclei. We identified pachytene nuclei by combining FISH and immunostaining of the SYCP3 axial component (Figure 2A, B and Figure S2A). The relative FISH area of the four tested autosomes was increased (from 1.3-fold to 1.6-fold) in *Ncapd2* cKO compared with WT samples (Figure 2C, D and Figure S2B, C). This indicates that the entire chromatin of the autosome bivalents 2, 5, 8 and 17 occupies a larger space in the absence of NCAPD2, independently of the chromosome size, suggesting that NCAPD2 promotes the global chromatin compaction of autosomes during prophase I.

Next, we asked whether the NCAPD2-dependent regulation of the global chromosome compaction correlated with a control of the local chromatin loop compaction. Previous analysis estimated that zygotene/pachytene loop size ranges from 250 to 500 kb ^10^. Thus, as a proxy of loop compaction, we measured the FISH signal extension from the axes of probes composed of three fosmids that cover contiguous portions of 121 kb and 126 kb on the subtelomeric regions of chromosomes 2 and 17 (Figure 2E, F). On both chromosomes, signal extension was significantly increased by 20% in *Ncapd2* cKO compared with WT samples (Figure 2G, H). This indicates that NCAPD2 controls chromatin compaction on autosomes by controlling loop extension.

Chromatin loop organization may not be evenly distributed in the genome. Particularly, the Pseudo Autosomal Region (PAR) of the X-Y chromosomes exhibits a disproportionately long axis relative to the DNA length and accordingly, smaller loops compared with autosomes ^41^. This organization correlates with higher DSB formation frequency. Therefore, we determined whether condensin I control chromatin compaction also in this specific region. Using whole chromosome X and Y FISH probes (Figure 2I, J), we measured a similar global chromatin occupancy for the two sex chromosomes in WT and *Ncapd2* cKO pachytene nuclei (Figure 2K, L). To restrict the analysis to the PAR region, we then used a combination of three fosmids at the PAR boundaries of the X chromosome (Figure S2D and Material and Methods). This approach showed that: (i) in WT samples, as already shown ^41^, the loop size was reduced in the PAR compared with autosomes (1.01 ± 0.4 µm vs 2.13 ± 0.9 µm and 2.40 ± 1 µm for autosomes 2 and 17, respectively), with a loop extension range similar between autosomes 2 and 17 (Figure S2E). And, (ii), as observed with the whole chromosome probes, signal extension with respect to the axis was similar in WT and *Ncapd2* cKO spermatocytes on sex chromosomes (Figure S2F). This suggests that chromatin compaction in the X-Y chromosomes, and in particular in the PAR region, is not driven by condensin I activity.

### The recruitment of the SYCP3 and cohesin axis components is NCAPD2- independent

Due to the close relationship between chromatin organization and the SC axial element, we investigated whether condensin I controls the organization and recruitment of major axial proteins: the SYCP2/SYCP3 structural meiotic proteins, the cohesin complexes, and HORMAD1/2.

In yeast and nematodes, the axis length is limited by the condensin I complex, which may influence chromatin loop organization ^15,27,33^. In WT mouse spermatocytes, SYCP3 assembles in a complex with SYCP2, and forms stretches that mark the axes (Figure 3A) ^11^. Thus, we used SYCP3 to measure the total axis length per nucleus on chromosome spreads. Axis length was similar in WT and *Ncapd2* cKO spermatocytes: 311 ± 40 µm vs 310 ± 38 µm at leptonema and 179 ± 24 µm vs 175 ± 30 µm at pachynema (Figure S3A). To exclude the possibility that mild differences on specific chromosomes may be undetectable with this approach, we used the FISH data performed on chromosomes 2, 5, 8, 17, X and Y (Figure 2 and Figure S2) to measure the axis length of chromosomes with different sizes, at pachynema. Again, we did not detect any significant difference in the axis length in the absence of NCAPD2 (Figure S3B). These results indicate that SYCP3 recruitment and axis length are NCAPD2-independent in the mouse. This conclusion was reinforced by the similar SYCP3 signal intensity per nucleus in WT and *Ncapd2* cKO spermatocytes (Figure S3C). As SYCP3 staining was used to identify the prophase I substage in all the experiments and because its intensity was similar in WT and *Ncapd2* cKO mutants, we decided to use SYCP3 total signal intensity per nucleus as a normalization factor for the other proteins, to compensate for potential experimental variation.

**Figure 3.**
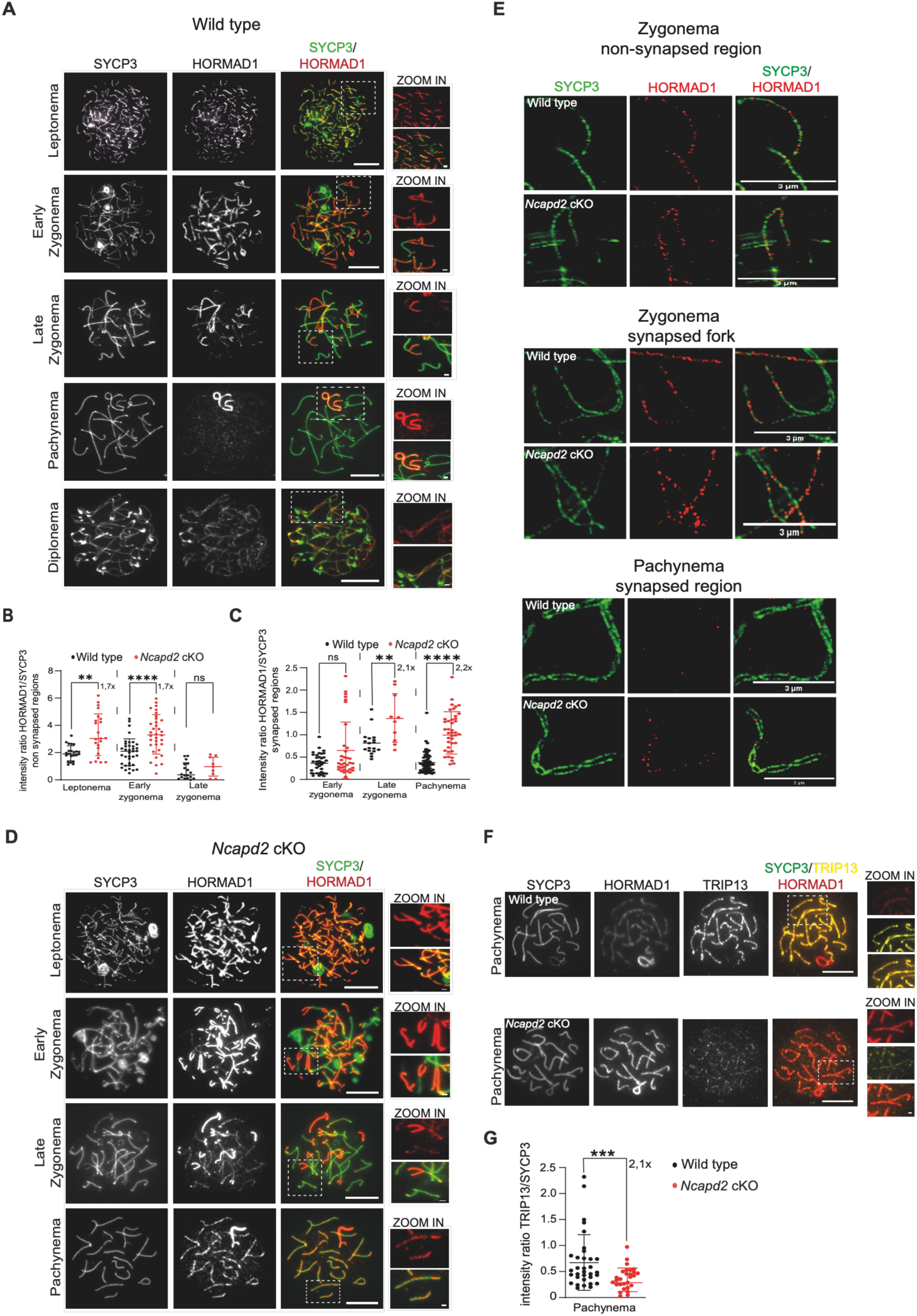
NCAPD2 controls HORMAD1 dynamics during prophase I. (A) Representative images of SYCP3 (green) and HORMAD1 (red) immunostaining from leptotene to diplotene nuclei in WT spermatocytes. (B) Quantification of HORMAD1 signal intensity on non-synapsed regions. Ratio of HORMAD1 signal intensity relative to SYCP3 on non-synapsed regions from leptonema to late zygonema in WT and *Ncapd2* cKO spermatocytes. Number of nuclei for WT and *Ncapd2* cKO, respectively, at leptonema: 23 and 23; early zygonema: 37 and 34; late zygonema: 18 and 10. (C) Quantification of HORMAD1 signal intensity on synapsed regions. Ratio of HORMAD1 signal intensity relative to SYCP3 on synapsed regions from early zygonema to pachynema in WT and *Ncapd2* cKO spermatocytes. The number of nuclei at early zygonema and late zygonema is the same as in (B), and at pachynema: WT: 44, *Ncapd2* cKO: 40. (D) Representative images of SYCP3 (green) and HORMAD1 (red) immunostaining from leptonema to pachynema in *Ncapd2* cKO spermatocytes. (E) HORMAD1 remains on the synapsed axes in *Ncapd2* cKO spermatocytes. Representative STED microscopy images of SYCP3 (green) and HORMAD1 (red) immunostaining in WT and *Ncapd2* cKO zygonema and pachynema nuclei. The three panels highlight the synapsed status of the chromosomes, from non-synapsed (*top)* to fully synapsed (*bottom*). Scale bar, 3 µm. (F) Representative images of SYCP3 (green), HORMAD1 (red) and TRIP13 (blue) immunostaining in pachytene WT and *Ncapd2* cKO nuclei. (G) TRIP13 loading depends on NCAPD2. Ratio of TRIP13 signal intensity relative to SYCP3 in WT and *Ncapd2* cKO pachytene nuclei. Number of nuclei at pachynema: WT: 33, *Ncapd2* cKO: 28. In all experiments, 18 dpp mice were used. White boxes indicate zoomed areas. Scale bars, 10 µm and 2 µm for zoomed images unless otherwise mentioned. P values were determined using the two-tailed unpaired Mann–Whitney test. Black and red bars show the mean values ± SD.

SMC-cohesin complexes also compose the axis ^10^. They hold together sister chromatids and are proposed to shape prophase I chromatin. In yeast, cohesin removal at the end of prophase I depends on condensin ^26^. In mammals, two meiotic specific *a*-kleisin subunits of cohesin have been identified, RAD21L and REC8, that define two different complexes. In WT leptotene spermatocytes, RAD21L and REC8 appeared as short stretches that colocalized with SYCP3 (Figure S3D). At zygonema, they formed segments along the axis that remained on the SC up to early pachynema (Figure S3D). At mid-pachynema, RAD21L signal started to dissociate from the SC, while REC8 remained associated with the axis up to diplonema (Figure S3D). In the absence of NCAPD2, RAD21L and REC8 recruitment patterns on the SC (Figure S3E) and signal intensity were unchanged compared with the WT condition (Figure S3F, G). This indicates that NCAPD2 is not required to control cohesin recruitment on the axis.

### NCAPD2 controls HORMAD1 pattern on the axes

The HORMAD1/2 HORMA domain-containing proteins are conserved axial components that coordinate HR and synapsis ^10^. In yeast, the proper recruitment of Hop1 (HORMA domain orthologue protein) depends on condensin ^27^ and in *Caenorhabditis elegans* the axis length increase observed in the condensin I mutant depends on HIM3 (HORMAD1 orthologue) ^33^. Therefore, we determined whether HORMAD1 recruitment in the mouse depends on NCAPD2 by monitoring its immunolocalization in WT and *Ncapd2* cKO spermatocytes.

In WT spermatocytes, HORMAD1 started to colocalize with SYCP3 along the forming axes at leptonema. During zygonema, it remained associated with the non-synapsed regions of the axes, while it was progressively removed from synapsed regions (Figure 3A-C). At pachynema, when chromosomes are fully synapsed, HORMAD1 became barely detectable on autosomes (Figure 3A- C). The increase in the signal intensity ratio within synapsed regions from early to late zygonema was consistent with synapsis progression, while the strong decrease at pachynema highlighted HORMAD active removal from synapsed chromosomes (Figure 3C). In WT pachytene nuclei, HORMAD1 remained abundant on the sex chromosomes, except in the synapsed PAR region (higher magnification in Figure 3A). Finally, in diplonema, when the SC is dismantled, HORMAD1 was reloaded onto the non-synapsed region of the axes (Figure 3A). At leptonema and early zygonema, in *Ncapd2* cKO spermatocytes, we observed excess HORMAD1 loading on non-synapsed regions, which is the main chromosome status at these stages (Figure 3D). In agreement, HORMAD1 relative intensity was increased by 1.7-fold compared with WT (Figure 3B). At late zygonema and pachynema, when synapsis is completed, HORMAD1 signal persisted on synapsed regions (Figure 3D), as indicated by the 2.1 to 2.2-fold increase of relative intensity compared with WT samples (Figure 3C). However, on the sex chromosomes at pachynema, HORMAD1 was present on the non-synapsed regions and absent in the PAR region in both mutant and WT samples. These results suggest excess HORMAD1 loading during axis formation on the autosomes, that accumulates or is only partially removed and persists upon synapsis in *Ncapd2* cKO spermatocytes. To thoroughly characterize HORMAD1 loading pattern in *Ncapd2* cKO spermatocytes and its localization relative to the axis and the SC central region, we used super-resolution stimulated emission depletion (STED) microscopy. In WT nuclei, at zygonema, non-synapsed regions exhibited multiple adjacent HORMAD1 foci that colocalized with SYCP3. At late zygonema and pachynema, HORMAD1 was not detectable on synapsed regions, in which SYCP3 was organized as two parallel lines, representing the two aligned lateral elements, sealed by the SC central element (Figure 3E). In *Ncapd2* cKO nuclei, despite the increased HORMAD1 relative signal intensity (quantification of the wide-field microscopy signal) (Figure 3B), we did not observe qualitative differences in HORMAD1 distribution on non-synapsed chromosomes by STED microscopy (Figure 3E). Conversely, on the synapsed regions at zygonema and pachynema, we detected discrete HORMAD1 foci that colocalized with the SYCP3 signal (Figure 3E). This confirmed that in the absence of NCAPD2, HORMAD1 persists on the SC upon synapsis, and that it remains associated with the SC lateral elements. Therefore, we propose that NCAPD2 is required to limit HORMAD1 loading on the axis and potentially to promote its removal during synapsis.

HORMAD1 removal from synapsed regions depends on the TRIP13 AAA+ ATPase ^19,42^. It was proposed that TRIP13 is recruited on the SC central element in pachytene nuclei to promote HORMAD1 removal through a HORMA domain-remodelling mechanism ^43,44^. TRIP13 immunostaining in spermatocytes showed that in WT pachytene nuclei, TRIP13 formed stretches that co-localized with the SC central region, as previously reported ^43^ (Figure 3F). In the absence of NCAPD2, TRIP13 relative intensity on the synapsed regions (TRIP13/SYCP3 intensity ratio) was reduced by ∼2.1-fold compared with WT nuclei (Figure 3G). This indicates that TRIP13 recruitment on the SC central element depends on NCAPD2, which correlates with HORMAD1 control.

### NCAPD2 regulates the SC central element assembly

As HORMAD1 removal and SYCP1 recruitment are related events ^19^, and considering the reduction in TRIP13 recruitment in the central region of the SC, we assessed whether NCAPD2 promotes the proper formation of the SC central region. First, we investigated the immunolocalization of three components of the central region: SYCP1 (transverse filament protein), TEX12 and SIX6OS1 (central element proteins) ^11^. In WT spermatocytes, these proteins started to be detected at zygonema, at the start of the central region formation (Figure 4A, B and S4A, E). In pachynema, they appeared as continuous lines that overlapped with the SYCP3 signal on the fully synapsed autosomes and in the PAR of the sex chromosomes. In diplonema, they were removed upon desynapsis, which corresponds to the dismantling of the SC central element, with the exception of the remaining connections between homologues that correspond to the future chiasmata (Figure 4A, B and Figure S4A). In *Ncapd2* cKO spermatocytes, we detected SYCP1, TEX12 and SIX6OS1 signals as continuous lines, like in WT nuclei (Figure 4C, D and Figure S4B). In addition, the SC central region length at pachynema was similar in WT and mutant spermatocytes (Figure S4C). We conclude that NCAPD2 absence does not lead to major synapsis defects, and that this protein does not control the SC length. Nevertheless, the quantification of the relative signal intensities of SYCP1, TEX12 and SIX6OS1 at pachynema revealed a decrease by 1.6-, 1.8- and 1.6-fold, respectively, in *Ncapd2* cKO compared with WT spermatocytes (Figure 4E, F and Figure S4D), indicating that SC transverse filament and central element components are less loaded in the absence of NCAPD2. As we recently proposed in collaboration with Matos’ group that the CHD1 chromatin remodeller is an interactor of the SC central element SYCE1 ^45^, we analysed also its localization to further document changes in the SC central region in the absence of NCAPD2. At pachynema, in WT spermatocytes, we observed CHD1 as stretches that colocalized with the synapsed axes (Figure 4G). In the absence of NCAPD2, the relative CHD1 signal intensity was reduced by 2.2-fold (Figure 4G, H). This confirmed that NCAPD2 absence globally impairs the SC central region protein recruitment and support the identification of CHD1 as a central region protein.

**Figure 4.**
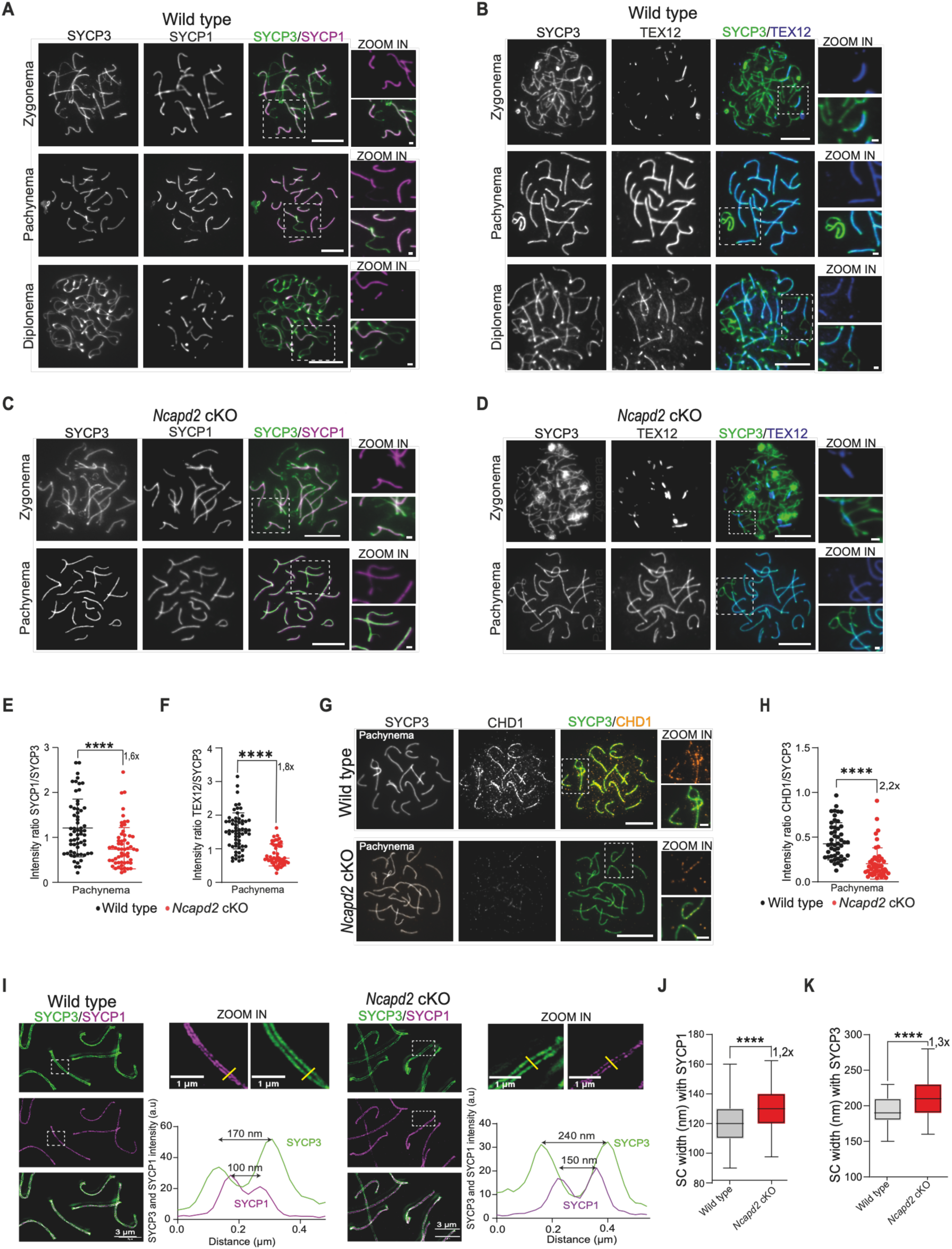
NCAPD2 controls the SC central element architecture. (A-D) SYCP1 and TEX12 loading. Representative images of SYCP3 (green) and SYCP1 (magenta) (A and C) and SYCP3 (green) and TEX12 (blue) (B and D) immunostaining from zygonema to diplonema (WT spermatocytes) and from zygonema to pachynema (*Ncapd2* cKO spermatocytes). (E and F) SYCP1 and TEX12 loading depends on NCAPD2. Ratio of SYCP1 signal intensity relative to SYCP3 in WT and *Ncapd2* cKO pachytene nuclei. Number of WT and *Ncapd2* cKO nuclei: SYCP1 staining: 61 and 57; TEX12 staining: 62 and 53. (G) Representative images of SYCP3 (green) and CHD1 (yellow) immunostaining in pachytene nuclei of WT and *Ncapd2* cKO spermatocytes. (H). CHD1 loading is NCAPD2-dependent. Ratio of CHD1 signal intensity relative to SYCP3 in WT and *Ncapd2* cKO pachynema spermatocytes. Number of nuclei at pachynema: 48 (WT), 50 (*Ncapd2* cKO). (I) SC width by STED microscopy. STED microscopy representative images of SYCP3 (green) and SYCP1 (magenta) immunostaining in WT and *Ncapd2* cKO pachynema. The yellow lines in the higher magnification image mark the SC width. SYCP3 and SYCP1 signal intensity profiles along the yellow line are plotted below the zoomed images. Scale bar: 3 µm and 1 µm for zoomed images. (J-K) SC width is regulated by NCAPD2. SC width quantification using SYCP1 (J) and SYCP3 (K) spacing in WT and *Ncapd2* cKO pachytene nuclei. SC width was measured randomly along the chromosomes and the number of measures is identical for SYCP1 and SYCP3 spacing: WT: 77; *Ncapd2* cKO: 85. All experiments were done with 18 dpp mice. White boxes indicate the zoomed images. Scale bars, 10 µm and 2 µm for zoomed images unless otherwise mentioned. P values were determined using the two-tailed unpaired Mann–Whitney test. Black and red bars show the mean values ± SD.

To better describe the NCAPD2-dependent control of the central element formation, we monitored SYCP1 and TEX12 distribution relative to SYCP3 by STED microscopy in pachytene nuclei. In WT, SYCP1 transverse filament appeared as two continuous linear lines, internal relative to the two SYCP3 axes, which corresponded to the staining of the SYCP1 polycomplex C-Terminal part (Figure 4I). We measured a regular spacing of the two SYCP3 axis lines and the two SYCP1 transverse filament lines at ∼190 ± 19 nm and 120 ± 16 nm, respectively, defining the SC width (Figure 4J-K and Materials and Methods section). In the absence of NCAPD2, the organization in continuous lines was maintained (Figure 4I), but the spacing increased to 212 ± 21 nm and 133 ± 13 nm for SYCP3 and SYCP1, respectively (Figure 4J-K). This indicates a 20%-30% increase of the SC width in the mutant condition.

In WT nuclei, the TEX12 central element protein appeared as a continuous line, running between the SYCP3 axes. Occasionally, this staining appeared as bubble-like structures, corresponding to regions where chromosomes twisted during spreading, as revealed by the SYCP3 signal (Figure S4E, white arrows). In *Ncapd2* cKO spermatocytes, TEX12 signal distribution was not altered, despite the reduction in the protein quantity measured after widefield microscopy imaging (Figure 4F).

The NCAPD2-dependent variation of the relative signal intensities of the SC central element proteins and HORMAD1 could be the consequence of transcription deregulation associated with condensin I depletion, as previously reported ^23^. The similar amounts of SC protein detected by western blotting in WT and *Ncapd2* cKO juvenile testis cells with synchronized spermatogenesis (Figure S4F) suggests that the observed phenotypes are not caused by transcription defects.

Altogether, we propose that the NCAPD2 condensin I subunit controls the global organization of the SC by (i) regulating HORMAD1 loading pattern, (ii) promoting the recruitment of SYCP1 transverse filament proteins and of central region proteins (TEX12, SIX6OS1 and CHD1), (iii) and limiting the SC width.

### NCAPD2 is not essential for DSB formation, but controls DMC1 loading

Meiotic HR takes place in the context of the structural reorganization of chromosomes and is affected by mutations in SC components. For example, DSB formation and processing are altered in *Hormad1^-/-^* mice ^16^, DSB repair and CO formation are altered in *Sycp3^-/-^* and *Sycp1^-/-^* mice ^46–48^, and DSB localization is affected by mutations that affect cohesin complexes ^49^. In yeast and *C. elegans*, it was proposed that the condensin I complex, by regulating axis length, directly controls DSB localization and level, thus affecting CO patterning ^27,28,33^.

To investigate whether NCAPD2 controls HR in mammals, we first monitored DSB formation through γH2AX localization ^50^. In WT spermatocytes, γH2AX was present as large chromatin domains that decorate nuclei at leptonema and zygonema, before its progressive relocalization on sex chromosomes at pachynema, forming the sex body (Figure S5A). In *Ncapd2* cKO nuclei, ɣH2AX presented similar kinetics and global intensity at prophase I onset (Figure S5A-B), showing that DSBs are formed in the absence of NCAPD2. However, at pachynema, in *Ncapd2* cKO nuclei, besides the normal accumulation in the sex body, discrete ɣH2AX foci accumulated on synapsed autosomes (Figure 5A). Compared with WT nuclei, ɣH2AX foci on synapsed regions were increased by 1.7-, 2.9- and 3.9-fold at early, mid and late pachynema, respectively (Figure 5B). This could be the consequence of a SC formation defect ^50,51^, but also the hallmark of a DSB repair defect, or an increase in their formation. We decided to explore these two last possibilities.

**Figure 5.**
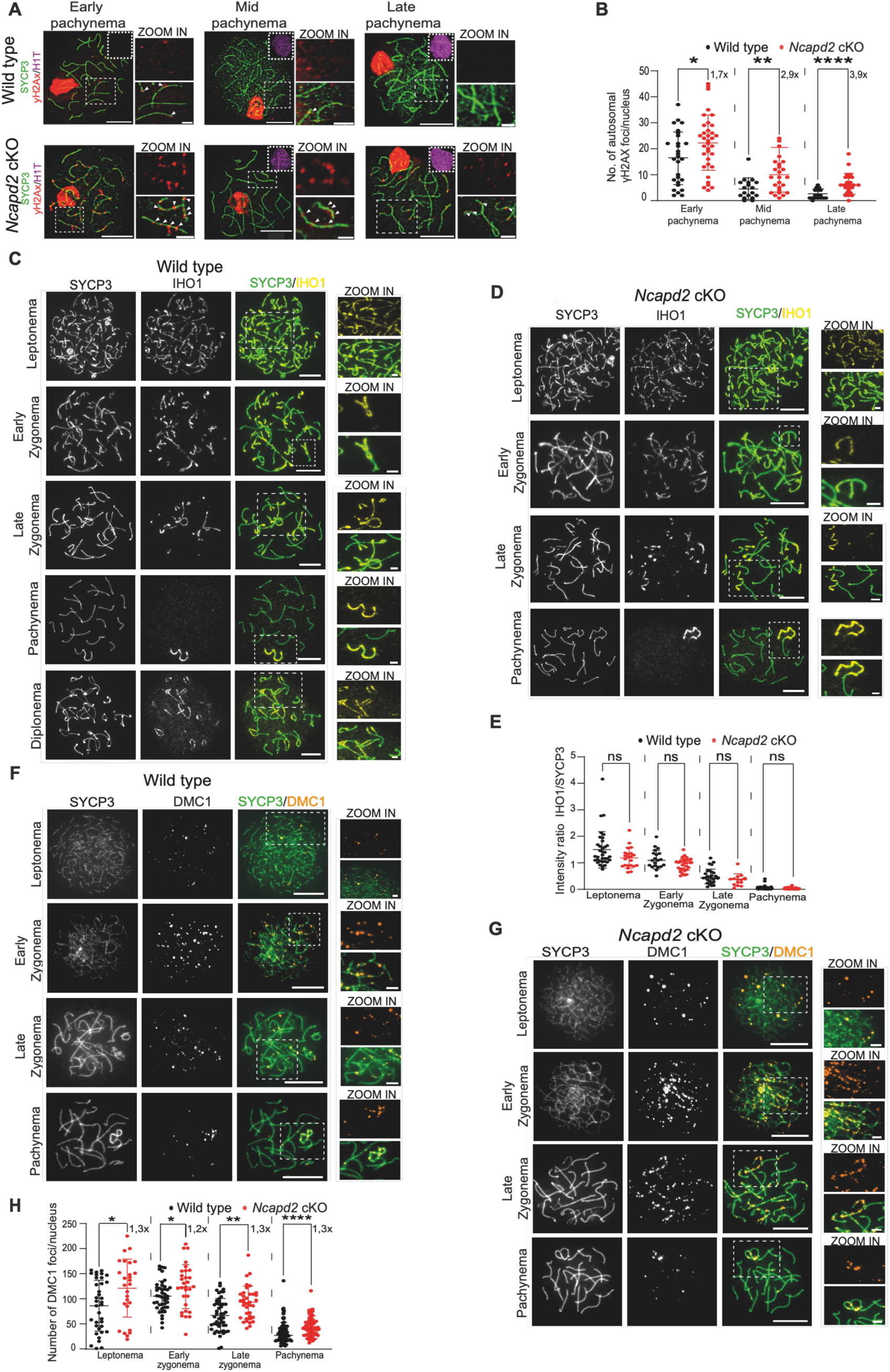
NCAPD2 controls γH2AX and DMC1 patterns. (A) Analysis of γH2AX localization in pachytene nuclei. Representative images of SYCP3 (green), γH2AX (red) and H1T (magenta) immunostaining in WT and *Ncapd2* cKO nuclei from early to late pachynema. White arrows mark γH2AX foci. (B) NCAPD2 promotes γH2AX focus removal at pachynema. Quantification of the number of autosomal γH2AX foci per nucleus in early, mid and late pachynema. Number of nuclei in WT and *Ncapd2* cKO nuclei at early pachynema: 28, 35; mid pachynema: 18, 25; and late pachynema: 17, 25, respectively. (C-D) Representative images of SYCP3 (green) and IHO1 (yellow) expression in WT and *Ncapd2* cKO nuclei. (E) IHO1 loading is NCAPD2-independent. Ratio of IHO1 signal intensity relative to SYCP3 from leptotene to pachytene nuclei in WT and *Ncapd2* cKO spermatocytes. Number of nuclei from WT and *Ncapd2* cKO spermatocytes at leptonema: 35, 22; early zygonema: 22, 29; late zygonema: 28, 15; and pachynema: 90, 72, respectively. (F-G) Representative images of SYCP3 (green) and DMC1 (orange) expression in WT and *Ncapd2* cKO spermatocytes. (H) DMC1 loading is NCAPD2-dependent. Number of DMC1 foci from leptonema to pachynema in WT and *Ncapd2* cKO spermatocyte nuclei. Number of WT and *Ncapd2* cKO nuclei at leptonema: 36, 27; early zygonema: 43, 30; late zygonema: 51, 37; and pachynema: 80, 68, respectively. For all experiments, 18 dpp mice were used. White boxes indicate the zoomed images. Scale bars, 10 µm and 2 µm for zoomed images. P values were determined using the two-tailed unpaired Mann–Whitney test. Black and red bars show the mean values ± SD.

In the mouse, meiotic DSB formation depends on the SPO11-TOPOVIBL core complex ^4,52^ assisted by its accessory factors REC114, MEI4 and IHO1 (RMI complex) and MEI1 ^3,53^. The three proteins of the RMI complex co-localize with the axis at leptonema and zygonema, independently of DSB formation ^3^. It was proposed that through a direct interaction with HORMAD1, IHO1 interacts with the axis and then recruits the REC114-MEI4 subcomplex, via the direct IHO1-REC114 interaction, forming the RMI complex ^8,53–56^. IHO1 may then act as an axis anchor that links the axis proteins with the DSB formation proteins. In line with this, HORMAD1 displacement upon synapsis might lead to the RMI complex removal and DSB formation activity downregulation. To determine whether NCAPD2, which controls HORMAD1 localization pattern, influences RMI recruitment and DSB formation, we monitored IHO1 and MEI4 localization. In WT nuclei, we detected the highest IHO1 signal as stretches that colocalized with the axes at leptonema. Then, this signal decreased at zygonema upon synapsis and HORMAD1 removal and completely disappeared at pachynema on autosomes. At this stage, IHO1 persisted only on the unsynapsed axis of the sex chromosomes (Figure 5C). MEI4 signal exhibited a similar kinetic, but it appeared as foci: 103.4 ± 63 in leptonema, 41 ± 28 in early zygonema upon synapsis formation, and none in pachynema (Figure S5C, D). In *Ncapd2* cKO nuclei, IHO1 and MEI4 patterns and signal intensity/focus number were similar to what observed in WT nuclei (Figure 5D, E and Figure S5C, D). We conclude that the recruitment of DSB formation proteins is NCAPD2-independent.

We next monitored DSB-end processing and invasion steps by immunolocalizing the DMC1 strand exchange protein. DMC1, the meiosis-specific paralogue of RAD51, assembles onto resected DSB ends to form nucleoprotein filaments that catalyse the homology search and strand exchange reaction ^1^. In WT nuclei, DMC1 appeared as foci along the chromosome axes during leptonema that persisted on both synapsed and non-synapsed regions through zygonema, and then disappeared at pachynema (Figure 5F). In the absence of NCAPD2, the number of DMC1 foci was increased by 20% to 30% from leptonema to pachynema (Figure 5G, H). This suggests that NCAPD2 limits DMC1 loading and/or potentially promotes its removal. However, we cannot distinguish whether it exerts this activity by limiting DSB level, controlling their repair kinetic, or enabling their final repair.

### NCAPD2 controls the maturation of the late recombination intermediates

Lastly, we wanted to determine whether NCAPD2 is required for the proper maturation of late recombination intermediates, as described in yeast and *C. elegans* ^27,33^. Following strand invasion of the homologous duplex, D-loops are formed, a subset of which will be converted into dHJ, and finally resolved as COs. In the mouse, 90% of COs are catalysed by the MutLɣ endonuclease complex (MLH1/MLH3), which defines the class I CO pathway. In WT spermatocytes, MLH1 formed one-two discrete foci per bivalent at mid- and late-pachynema that mark the future CO sites and the chiasmata (Figure 6A). In the absence of NCAPD2, we found: (i) a delay in MLH1 focus formation, which started to appear at late pachynema, and (ii) a slight increased by 10% in MLH1 foci number at late pachynema (Figure 6A-B).

**Figure 6.**
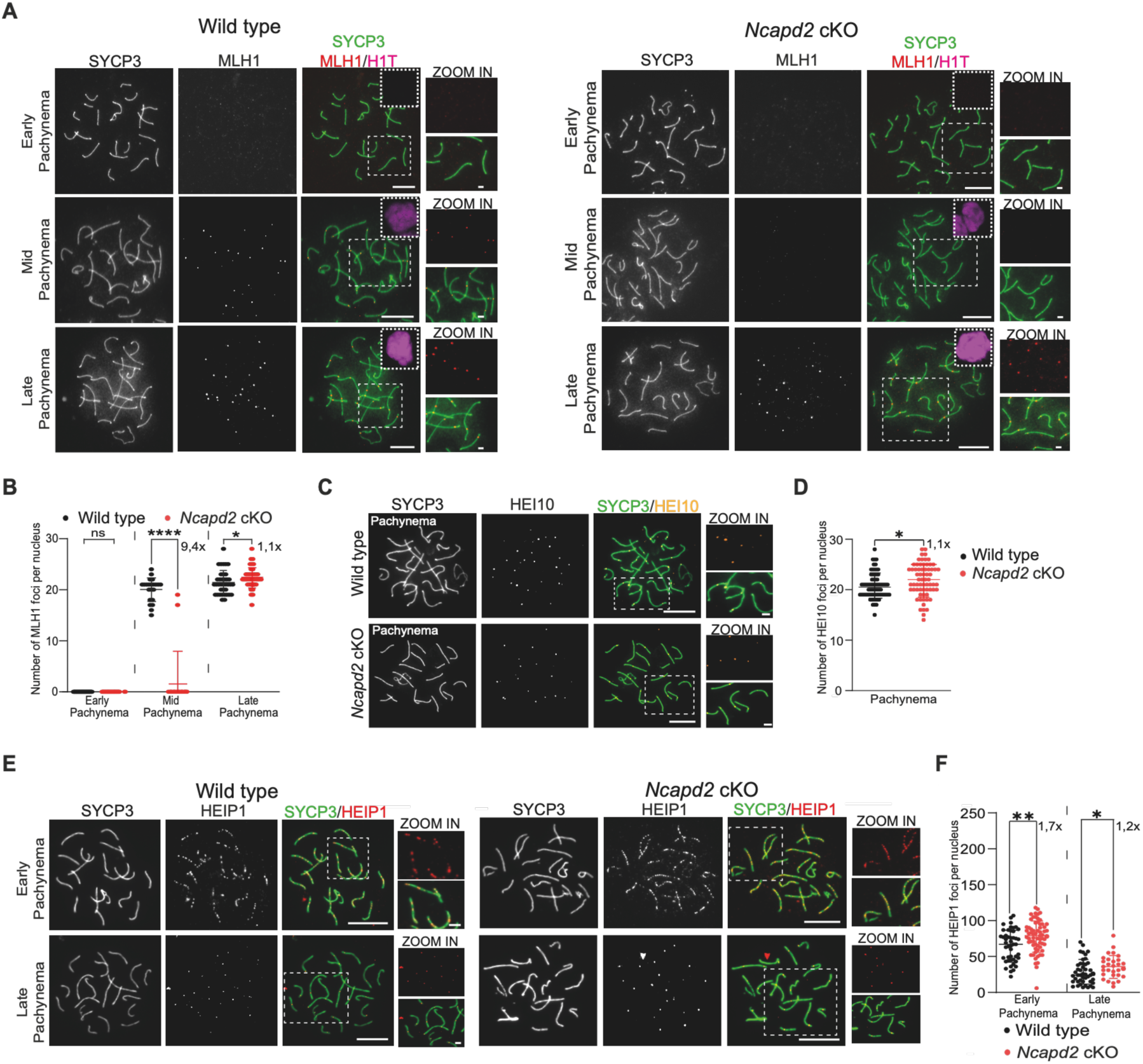
NCAPD2 controls crossovers and non-crossover protein loading. (A) MLH1 loading in WT and *Ncapd2* cKO. Representative images of SYCP3 (green), MLH1 (red) and H1T (magenta) immunostaining from early to late pachynema. (B) Number of MLH1 foci in WT and *Ncapd2* cKO spermatocyte nuclei from early to late pachytene. Number of WT and *Ncapd2* cKO nuclei at early pachynema: 45, 36; mid pachynema: 22, 25; and late pachynema: 37,38, respectively. (C) HEI10 loading in WT and *Ncapd2* cKO spermatocytes. Representative images of SYCP3 (green) and HEI10 (orange) immunostaining in pachytene nuclei of WT and *Ncapd2* cKO mice. (D) Number of HEI10 foci in pachytene nuclei of WT and *Ncapd2* cKO mice. Number of nuclei: 72 (WT), 66 (*Ncapd2* cKO). (E) HEIP1 loading in WT and *Ncapd2* cKO. Representative images of SYCP3 (green) and HEIP1 (red) immunostaining in early and late pachytene nuclei. (F) Number of HEIP1 foci in WT and *Ncapd2* cKO spermatocyte nuclei from zygonema to late pachynema. Number of WT and *Ncapd2* cKO nuclei at early pachynema: 47, 65; and late pachynema: 45, 29, respectively. All experiments were done using 18 dpp mice. White boxes indicate zoomed areas. Scale bars, 10 µm and 2 µm for zoomed images. P values were determined using the two-tailed unpaired Mann–Whitney test. Black and red bars show the mean values ± SD.

Class I CO formation depends on a set of conserved factors, the pro-crossover proteins, which progressively stabilize and direct dHJ resolution toward a proper number of COs ^5^. To evaluate whether condensin I is required for the general control of pro-crossover factors, we monitored the recruitment of HEI10 RING domain protein ^57^ and HEIP1 pro-crossover master regulator ^58^. In WT spermatocytes, HEI10 formed 21 ± 2 discrete foci from early to late pachynema that co-localized with MLH1, marking CO sites (Figure 6C, D). HEIP1 was loaded earlier on the chromosomes. It appeared as punctate foci at early zygonema that increased up to late zygonema, before their progressive reduction and relocalization to CO sites during pachynema (Figure 6E). As observed for MLH1, the number of HEI10 foci at late pachynema and of HEIP1 foci throughout pachynema was increased by 10% in the absence of NCAPD2 (Figure 6D, F).

Western blotting of whole protein extracts from WT and *Ncapd2* cKO juvenile testes cells with synchronized spermatogenesis showed similar amounts of the different studied HR proteins in both conditions (Figure S5E), indicating that the variations in the number or intensity of recombination protein foci in *Ncapd2* cKO nuclei were not related to a protein expression deregulation.

Altogether, we propose that NCAPD2 controls recombination progression and outcome by limiting the loading of pro-crossover proteins.

## Discussion

In mammalian meiosis, the identification and characterization of the factors that control the higher- order chromosome organization during prophase I remains an essential goal to fully understand the different layers of regulation of recombination and chromosome segregation. The factors that specifically drive chromatin folding remain poorly known. Here, we extensively characterized the SMC-condensin I complex role in these processes and found that it acts as a prophase I chromatin organizer that regulates SC assembly and HR outcome.

### Condensin I: a driver of chromatin compaction during mouse prophase I

SMC-condensin complexes are one of the main drivers of chromatin organization. Different studies showed that condensin I and II collaborate in the assembly of the vertebrate mitotic chromosome ^21,22,59^. It was proposed that condensin II controls the chromatid longitudinal compaction by initiating the formation of chromatin loops, followed by a higher-order assembly that leads to a linear organization and axial control of chromosome length. Then, condensin I could contribute to the chromosome lateral compaction by further reorganizing the initial chromatin loops through the formation of smaller nested loops ^21,24^ In that scenario, condensin I and II roles rely on a loop formation activity, potentially mediated by a loop extrusion mechanism ^25,60,61^. In mammalian meiosis, the function of condensin was documented in metaphase, with a more pronounced role for condensin II, which promotes the chromosome longitudinal rigidity ^62^. In our study, we show that in prophase I, condensin I limits autosomes chromatin occupancy by controlling loops organization. We obtained these results in four different autosomes (2, 5, 8 and 17), the size of which ranges from ∼95 Mb to ∼181 Mb, thus covering the spectrum of mouse chromosome sizes. We conclude that this effect is common for all the autosomes and not dependent on their size. Then, by using fosmid FISH probes that cover contiguous regions of ∼125 Kb, we found that NCAPD2 limits chromatin extension from the axis. Therefore, we propose that condensin I promotes chromatin compaction by constraining the prophase I chromatin loop size. Based on the observation that loop density remains constant, the most common model postulates that prophase I chromatin loop size is controlled by the length of the chromosome axis on which they are anchored: a long axis leads to shorter loops ^40^. In our assay, the length of the chromosome axis was condensin I-independent (Figure S3 and below for discussion on SC architecture controls by condensin). Then, similarly to what has been postulated in mitosis, we hypothesise that in prophase I, condensin I promotes the formation of nested chromatin loops ^21,24^ in the context of the SC that favour the chromosome lateral compaction. This is in accordance with the chromatin localization of NCAPD2 (Figure 1), and provides a role for condensin I in promoting an additional layer of regulation of prophase I chromatin organization. Spatial genomic analyses and genome-wide mapping of condensin I binding sites are needed to further validate and document this model.

In our assay, in the X-Y sex chromosomes, particularly in the PAR region, condensin I does not control chromatin compaction. This region of the genome is under a specialized regulation, which has important functional consequences. The PAR axis is longer than in the rest of the genome relative to the DNA length. This promotes the formation of shorter chromatin loops and increases the rate of DSB formation ^7,41^. We thus suggest that the chromatin organization in the PAR is mainly driven by an axis-specific regulation which promote the formation of smaller loops than in the rest of the genome that escape the condensin I control and is not prone to nested chromatin loop formation. In addition, the whole X chromosome FISH data suggest that the regulation on chromatin organization observed in the PAR extends on the rest of the chromosome. However, a more precise characterization of the PAR *vs* non-PAR regions remains to be done to validate this model.

### NCAPD2 controls HORMAD1 turnover, central element protein recruitment and SC width

Condensin’s role in controlling SC assembly in prophase I has been mainly documented in yeast and *C. elegans.* This is supported by the colocalization of different condensin subunits with the axis ^27,34,63^ and by the accumulation of the yeast complex in the vicinity of CO-designated sites on the SC^30^. Based on NCAPD2 immunodetection, we did not observe this localization pattern, but rather a chromatin-spread localization. This supports our model of a role for condensin I in controlling global chromatin folding. However, we do not exclude a local/transient condensin I enrichment at some specific axis/SC sites that our assay could not detect. In yeast and nematodes, condensin limits the axis length ^27,28,32,33^. In the mouse, using SYCP3 total measurement length per nucleus, as well as on four specific chromosomes, as a proxy, we did not detect a role for NCAPD2 in controlling the axis length. However, SYCP3 immunofluorescence-based assays may not be sensitive enough to detect subtle or local changes.

At the protein level, yeast Hop1, Red1 and Zip1 (respectively HORMAD1, SYCP2 and SYCP1) recruitment depends on condensin, as well as cohesin removal at prophase I end ^26,27^. In *C. elegans, Dpy-28* genetically interacts with *Him-3* (the *Hormad1* ortholog), and the cohesin pattern as well as SYP-1 (the SYCP1 orthologue) recruitment depends on condensin I. In the mouse, we also found that NCAPD2 limits HORMAD1 axial protein recruitment, and promotes its removal upon synapsis. In addition, we showed that proper central element protein recruitment requires NCAPD2. Altogether, it appears that the recruitment and assembly of axis/SC proteins controlled by condensin is a conserved feature. In the mouse in particular, HORMA domain components and central element proteins appear as more specific targets of condensin I regulation.

So far, a unified mode of action of condensin I in SC formation control is difficult to propose. One simple hypothesis is that this complex directly interacts with and controls the recruitment of some axis/SC proteins. Non-exclusively, and compatible with the condensin I chromatin localization that we reported, we propose that condensin I-dependent chromatin organization could promote the proper recruitment/removal of different axis/SC components, particularly HORMAD1 and central element proteins. An important quantity of DNA is embedded in the SC, chromatin is anchored on the axis and some of the proteins that compose it directly interact with DNA ^11,16,40^. It is thus tempting to propose that chromatin architecture drives axis/SC organization. Remarkably, we showed that NCAPD2 controls the proper recruitment of CHD1, a chromatin remodeller that interacts with SYCE1 and proposed to control prophase I chromatin organisation and SC assembly ^45^. This result confirms that NCAPD2 regulates central element components recruitment. But it also highlights the importance and potential inter-relationship of chromatin organization proteins in mouse prophase I. At the molecular level, NCAPD2 depletion, by decompacting chromatin, could modify the axis physical properties (*e.g.* axis protein rigidity could be altered), which then could modify the axial accessibility of proteins and lead, for example, to HORMAD1 accumulation. One possibility is that HORMAD1 axial element is more sensitive to these changes due to its dynamic pattern and connection with HR in prophase I. In turn, HORMAD1 accumulation could block the further recruitment of central element proteins that actually promote its own removal, particularly TRIP13 (Chotiner et al., 2024; Roig et al., 2010; Ye et al., 2017). This hypothesis is in line with different studies showing that HORMAD1 accumulation could hinder its structural reorganization, essential to its removal by TRIP13 ^18,19,44^. Alternatively, the change in SC central structure could specifically block TRIP13 recruitment in pachynema.

Lastly, we found that condensin I controls the global super-structure of the SC in pachynema by limiting its width all along the synapsed chromosomes. The SC width is highly conserved and fixed (∼ 100 nm in mammals), thus implying a fine tuning activity to control its architecture ^11,40^. We propose that the SC width regulation could be mediated by the chromatin packaging induced by condensin I. For instance, limiting chromatin occupancy in the central region of the SC or promoting lateral condensation of loops anchored on the axial/lateral element could limit SC spacing. Non-exclusively, this function could also be correlated to the SC protein organization mediated by condensin. The proper recruitment of HORMAD1 and central element proteins could be essential to drive the formation of a properly spaced SC. We did not detect defects in the homologue alignment and synapsis in the absence of NCAPD2 associated with the lack of SC width control. Nevertheless, we do not exclude the possibility of local and transient undetectable effects.

### Condensin I and meiotic HR control

Meiotic HR takes place in the context of the SC and of the specific loop/axis chromatin organization, highlighting their tight relationship. For example, knowing that DSBs are formed in the loops and repaired on the axis, chromatin compaction and loop/axis organization could control DSB number, location, timing of formation and repair. CO resolution is also directly connected with SC formation: pro-crossover proteins are present in the central part of the SC and the SC structure might directly influence their relocalization and accumulation on few foci that are the future CO sites ^5^.

We found that in the mouse, NCAPD2 controls recombination in prophase I: it promotes an efficient timing of repair and limits the accumulation of strand invasion-prone recombination intermediates (DMC1 protein) and late CO recombination intermediates (HEI10, HEIP1 and MLH1 proteins). This recombination regulatory effect is conserved in yeast where condensin controls CO outcome at specific hotspots ^30^, and nematodes where DSB and CO formation is regulated by condensin dependent axis length regulation ^33^. In the mouse, the mode of action of condensin remains elusive.

In particular, whether DSB formation is controlled by NCAPD2 remains unclear. We used ɣH2AX signal intensity as a proxy for DSB formation, but subtle differences in DSB numbers and modification in their location are undetectable using this approach. The condensin I control of chromatin loop organization suggests that condensin I could control not only DSB level, but also their distribution in the genome.

Moreover, we propose that the condensin I chromatin organization activity could also promote efficient DSB repair: recombination sites properly regulated in number and location will be efficiently repaired by HR. However, we envisage a non-exclusive level of regulation mediated by the control of HORMAD1 recruitment and its link with the meiotic checkpoint ^18,19^. Indeed, HORMAD1 accumulation in *Spo11^-/-^* spermatocytes promotes ATR hyper-recruitment and activity, which could promote ɣH2AX accumulation, associated with synapsis defect. In this context, we could hypothesize that condensin I, by controlling HORMAD1 removal, could contribute to ATR regulation. This is in line with the delay in prophase I progression in the absence of NCAPD2, but not consistent with the absence of synapsis defects. Lastly, in line with a potential increase/reshuffling of DSB formation or recombination delay in the absence of NCAPD2, we found a slight delay and increase in the level of late pro-CO recombination markers. Such increase was also observed in yeast, at specific CO locations ^30^, and in nematodes ^33^.

Altogether, we think that condensin I-dependent control of chromatin organization is required to promote proper recombination, up to the resolution of late recombination intermediates. Finally, in addition to impacting HR from the early steps, we propose two non-exclusive hypotheses to account for the NCAPD2 specific CO control: (i) chromatin reorganization could influence the late DNA recombination intermediate structure and then their resolution; and (ii) SC width control could contribute to the regulation of pro-CO proteins.

### Conclusion and model for condensin I activity in prophase I

Based on our results on the NCAPD2 SMC-condensin I subunit, we propose that condensin I triggers the higher-order chromosome organization in prophase I through the control of chromatin 3D architecture and synaptonemal complex structure. This has a direct effect on HR. We propose the following dynamic model (Figure 7). Condensin I first localizes on the forming chromatin loops that emanate from the SC throughout prophase I (Figure 7 (1)). Then, condensin I catalyse the formation of nested loops throughout the genom, promoting a higher level of chromatin compaction (Figure 7 (2)). We propose that this activity, by constraining the axis physical properties, regulates HORMAD1 loading and turn-over on the axis (Figure 7 (3)). This condensin I activity could then promote the assembly of an equilibrated synapsed SC with a 100 nm SYCP1 spacing (Figure 7 (4)). Finally, this chromatin organization could play a direct role in controlling DSB localization (*e.g.* by influencing DSB site location) and, timely and efficient repair by recombination (Figure 7 (5)).

**Figure 7.**
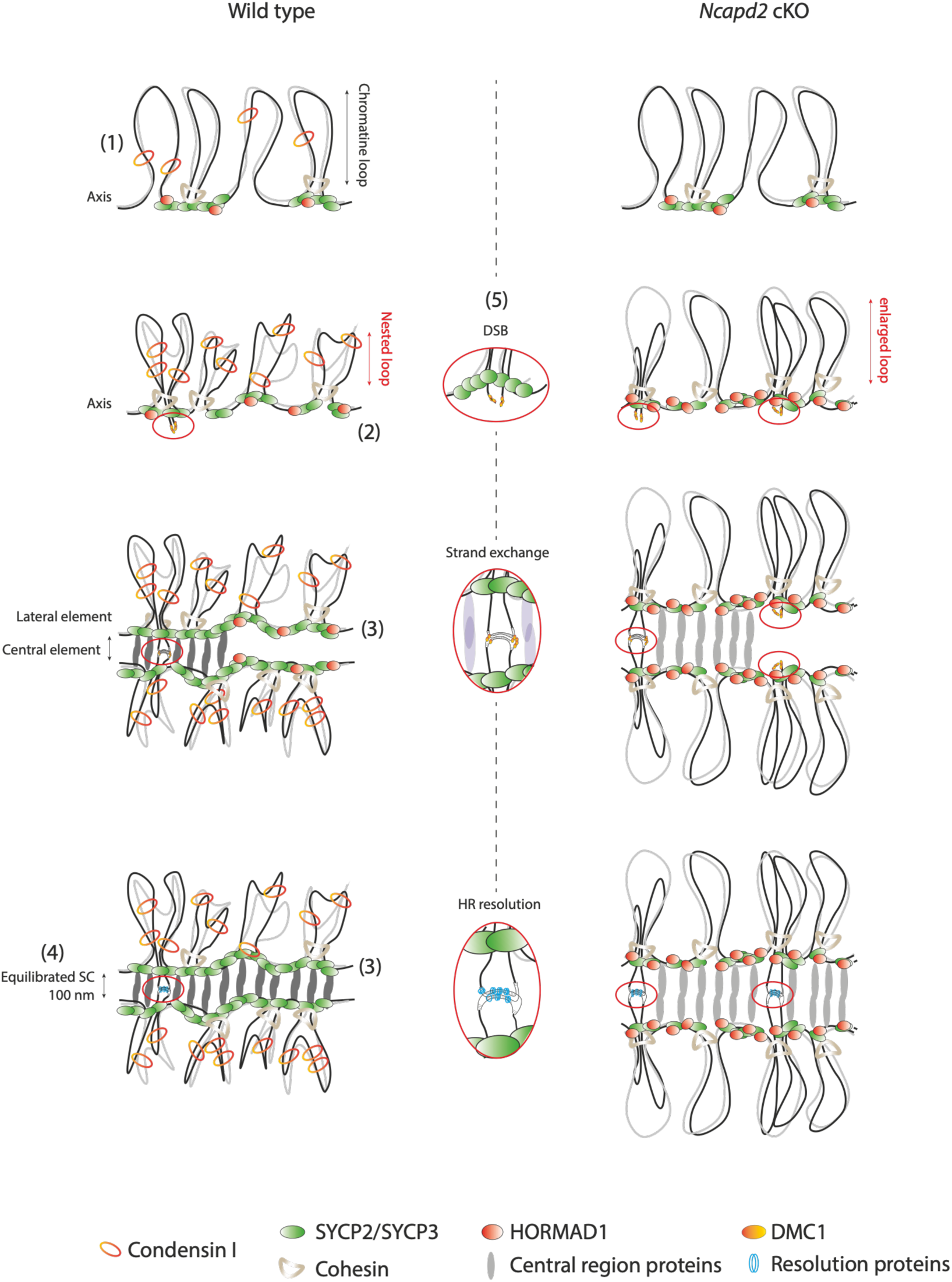
Model for NCAPD2 role in prophase I chromosome organization. (*Left)* Condensin I mode of action in WT mice. (1) Condensin I is loaded on chromatin at the onset of prophase I, concomitantly with axis formation. (2) During chromosome organization, condensin I catalyses the formation of nested loops. (3) This chromatin folding influences the axis physical structure, thus promoting the proper turn-over of HORMAD1 and central element loading and leading to the formation of a properly spaced SC (4). (5) Condensin I chromatin activity controls HR outcome. *(Right*) In the absence of NCAPD2, nested loops do not form and chromatin is more extended. In turn, HORMAD1 accumulates on the axis, central element proteins are less loaded and the SC is enlarged. Consequently, HR intermediates accumulate.

## Materials and Methods

### Mice strains

All mice used in the study were in the C57BL/6J background. *Ncapd2^tm1a(EUCOMM)Wtsi^*embryonic stem cells were obtained from Helmholtz Zentrum München GmbH. *Ncapd2^tm1a(EUCOMM)Wtsi^* mice were generated by Phenomin-Institut Clinique de la Souris and mated with mice that express Flp under the control of the CMV promoter to generate *Ncapd2^tm1c^* (*Ncapd2^f/+^*)*. Ncapd2^flox/+^* mice were mated with Tg(Stra8-icre)1Reb/J *(Stra8-Cre^Tg^*) mice to generate *Ncapd2^f+^*;*Stra8-Cre^Tg^* mice. Then, *Ncapd2^f/+^*;*Stra8-Cre^Tg^* males were mated with B6 wild type females to generate *Ncapd2^Δ/+^*;*Stra8- Cre^Tg^* mice (Figure S1). *Ncapd2^flox/+^* mice were mated together to generate *Ncapd2^flox/flox^* mice. By crossing *Ncapd2^flox/flox^* mice with *Ncapd2^Δ/+^*;*Stra8-Cre^Tg^* mice, *Ncapd2^flox/Δ^*;*Stra8-Cre^Tg^* (*Ncapd2* cKO) and *Ncapd2^flox/+^* (*Ncapd2* control) mice were obtained. The following primers were used for genotyping: primer 361 (TGTGGTTCTGGGAATCAGCCC), primer 390 (GGTGAAGCTGGTGCATGGTGC) and primer 392 (CAGTGTGCCTATCTCCTCTGC). See Figure S1 for crossings and genotyping. All animal experiments were carried out according to the CNRS guidelines.

### Histological analysis of paraffin sections and TUNEL assay

Mouse testes were fixed in 4% paraformaldehyde/1X PBS for haematoxylin-eosin staining, immunostaining and TUNEL assay. Testes were embedded in paraffin and cut in 3 μm-thick sections. Sections were scanned using the automated tissue slide-scanning tool of a Hamamatsu NanoZoomer Digital Pathology system. TUNEL assays were performed with the DeadEnd Fluorometric TUNEL System (Promega), according to the manufacturer’s protocol.

### Spermatozoid counting

After dissection of the epididymis caudal part from adult (2-month-old) testes, spermatozoids were extracted from the epididymis by smashing or crushing the tissue in PBS. After homogenization by pipetting, 10 μl of the soluble part was diluted in 1mL of water, and spermatozoids were counted.

### Antibodies

The homemade polyclonal anti-NCAPD2 antibody was produced in goats. Two *Ncapd2* domains were amplified from mouse testis cDNA: Dom1 using the primers Oli307 (CGCGCTAGCATGCACCATCACCATCACCATCACCATCACCATAGTGCTTCTCCCCACAACTTT GAATTCC) and Oli308 (CCGCGGATCCTTAAGTGGCACTGAGCATATGGTTGTAG) and Dom2 using the primers Oli309 (GGCCATATGATCTGTGAGAAGGAGCTGTTGGACG) and Oli310 (GGCGCTAGCTTAATGGTGATGGTGATGGTGATGGTGATGGTGAGCACTGGACCTGTGTCTTCGCCCAGATGCTC). The obtained amplicons were cloned in pET11a at the NheI/BamH1 and NdeI/NheI sites for Dom1 and Dom2, respectively. Transformed *E. coli* (DE3) cells were grown in 2 L of LB at 37°C to OD_600_ = 0.8, and protein expression was induced with 0.5 mM IPTG followed by incubation at 37°C for 3h. Protein fragments were prepared from inclusion bodies, and co-injected (5 mg of each fragment) in goats. Goat serum was affinity-purified using both fragments separately, and subsequently mixed. The different antibodies used in the study are listed in Table S1.

### Immunocytology

Spermatocyte chromosome spreads were prepared with the dry down technique, as described in ^64^. Briefly, a testis cell suspension was prepared in PBS, and then incubated in hypotonic solution at room temperature for 8min. Cells were centrifuged, resuspended in 66 mM sucrose solution and spread on slides with 1% paraformaldehyde/0.05% Triton X-100. Slides were dried in a humid chamber for 1 to 2h. Immunostaining was performed using a milk-based blocking buffer (5% milk, 5% donkey serum in PBS) as in ^65^. Briefly, slides were incubated with primary antibodies at room temperature overnight and with secondary antibodies at 37 °C for 1h. Nuclei were stained with 2 μg/ml 4’−6-diamidino-2-phenylindole (DAPI) during the final washing step.

### Fluorescence In Situ Hybridization (FISH)

The FISH protocol was adapted from ^66^. Briefly, slides with chromosome spreads from spermatocytes were incubated with 1M NaSCN at 70°C for 30min, washed in 1X Saline Sodium Citrate (SSC) buffer, and digested with 100 µg/ml RNase A 37°C for 1h. Then, slides were washed in 1X SSC buffer and covered with 70% formamide/30% 2X SSC buffer, denatured by heating at 85°C for 5min, and washed in ice-cold 2X SSC to block renaturation. Probes in hybridization buffer (see below) were added on the slides and then covered with a cover slip, sealed with rubber cement and incubated in a humid chamber at 42°C for 12h. Then, slides were washed 3x5 min in 0.05X SSC buffer at 42°C and handled for immunostaining following standard procedure (see above).

Whole chromosome probes were obtained from Calibre Scientific: XMP2 D-1402-050-04, XMP5 D- 1405-050-0R, XMP8 D-1408-050-FI, XMP17 D-1417-050-FI. Probes (7 µL/each) were denatured at 75°C for 7min and then applied on the slides.

The fosmids used to measure the FISH signal extension were from BACPAC genomics: WIBR1- 2473G14, WIBR1-0132I08 and WIBR1-0744O09 (for chromosome 2); WIBR1-0501A19, WIBR1- 2526E06 and WIBR1-0119I02 (for chromosome 17); and WIBR1-1482I02, WIBR1-1993H01 and WIBRE1-1291N07 (for chromosome X). The X fosmids are located at the PAR boundary, with WIBR1-1482I02 (33.8 Kb) within the PAR and WIBR1-1993H01 (39 Kb), WIBRE1-1291N07 (40.1 Kb) X specific 1kb close to the PAR. Fosmids were amplified from bacterial cultures in LB medium supplemented with 25 µg/mL chloramphenicol and extracted using a maxi-prep kit (Macherey Nagel) according to the manufacturer’s protocol. For labelling, 1 µg of each fosmid was used. The three fosmids corresponding to each chromosome were labelled together with the FISH Tag™ DNA Multicolor Kit, Alexa Fluor™ dye combination (Invitrogen, F32951) according to the manufacturer’s protocol. For one slide, 40 ng of labelled probes, 10 µg of salmon sperm DNA (Invitrogen, 15632011) and 5 µg of mouse Cot-1 DNA (Invitrogen, 18440016) were precipitated with 200 µL of EtOH/0.096 M AcNa and then washed with 70% ethanol. The pellet was resuspended in 5 µL of 100% formamide at 37°C, probes were denatured at 75°C for 7min, and added to the slides at 37°C for 1h. Finally, 4 µL of 20X SSC buffer and 6µL of 50% dextran were added.

### Widefield fluorescent and stimulated emission depletion (STED) super-resolution imaging

Widefield images were acquired using an Axioimager Z2 Apotome (Zeiss) with 63X Plan Apochromat 1.46 NA oil objective and an ORCA-Flash4 LT (Hamamatsu) monochrome camera (2048 x 2048 pixels, 6.5 µm pixel size).

Super-resolution images were acquired using a Stimulated Emission Depletion (STED microscope, Expert Line, Abberior Instruments) equipped with a PlanSuperApo 100x/1.40 oil immersion objective (Olympus). For STED microscopy, the following secondary antibodies were used: anti-mouse STAR ORANGE, anti-rabbit STAR RED or anti-mouse STAR ORANGE, anti-guinea pig STAR RED (Table S1). STAR ORANGE was excited at 561 nm, and STAR RED at 640 nm. Excitation was done with a dwell time of 10 μs. STED was performed at 775 nm for STAR ORANGE and STAR RED. Images were collected in line accumulation mode with detection set at 580-630 nm for STAR ORANGE and 650-750 nm for STAR RED.

### Image analysis

Staging criteria were as follows. Leptotene nuclei had only short SYCP3 fragments, 100% of HORMAD1 staining colocalized with SYCP3, and no SYCP1 or TEX12 staining. Early zygonema had <50% of synapsed homologues (the percentage of synapsed homologues was calculated using the ratio of the SYCP1 or TEX12 length to the SYCP3 length). Late zygotene nuclei had >50% of synapsed homologues, and pachytene nuclei had all 19 autosomes fully synapsed. When stated, pachytene was divided in three sub-stages (early-, mid- and late-pachynema) based on H1T staining quantification: no/weak H1T intensity for early, H1T mid-intensity for mid, and strong H1T intensity for late pachynema.

FISH quantifications were done on non-deconvoluted images using the ImageJ software. The FISH probes area, FISH signal extension and DAPI area were measured using the freehand line tool. To normalize the FISH area measurement, the ratio between the FISH probe area and the corresponding DAPI area on the same nucleus was calculated.

For immunostaining quantification, images were deconvoluted using the Huygens Professional software and image analyses (foci count, intensity and length measurements) were performed using Fiji/ImageJ 1.53t and the “MeiQuant” set of tools ^67^ available on github (https://github.com/MontpellierRessourcesImagerie/meiosis_bar). Only, MLH1 and HEI10 foci were counted manually. All intensity measurements were normalized to the SYCP3 intensity. For γH2AX, MLH1 and HEI10 quantification, foci were counted manually on non-deconvoluted images.

The SC width was measured by drawing regularly spaced lines perpendicular to the SC, which correspond to SC section, on several chromosomes using ImageJ (10 to 15 per chromosome, depending on their size). For each section, the plot profile intensity was generated by ImageJ. Each graph in Figure 4I shows an example of one section intensity profile where the SC width (in nm) is defined as the distance between the centres of the two SYCP3 (or SYCP1) peak signals.

### Preparation of mouse protein extracts and western blotting

*Whole proteins extracts:* four testes per genotype were homogenized using a Dounce homogenizer in HNTG buffer (150 mM NaCl, 20 mM HEPES pH 7.5, 1% Triton X-100, 10% glycerol, 1 mM MgCl₂, Complete protease inhibitor [Roche 11873580001]), followed by sonication. Benzonase (250 U) was added at 4°C for 1h. After centrifugation (16000 g, 4°C, 10 min), the supernatant was collected (whole cell protein extract). *Nuclear extracts:* four testes were homogenized using a Dounce homogenizer in homogenizing buffer (10 mM HEPES pH 7.4, 320 mM sucrose, 1 mM PMSF, 1x protease inhibitor cocktail) and centrifuged at 1000 g at 4°C for 10 min. The supernatant was collected (cytoplasmic fraction). The pellet was resuspended in RIPA buffer (50 mM Tris–HCl pH 7.5, 150 mM NaCl, 1 mM EDTA, 1% NP-40, 0.5% sodium deoxycholate, 0.1% SDS, protease inhibitors) and then sonicated. Following centrifugation, 16000 g at 4°C for 10 min, the supernatant was collected (nuclear extract).

The expression of NCAPD2, SMC2, NCAPD3, HORMAD1, SYCP1, TEX12, TRIP13, MLH1, HEI10, DMC1 and β-tubulin in whole cell extracts, and of NCAPD2 and β-tubulin in nuclear extracts, were assessed by western blotting and the antibodies listed in Table S1.

### Statistical analysis

The statistical analyses of cytological observations were done with GraphPad Prism 10. The one-sample t-test or the nonparametric Mann-Whitney test was used to compare signal intensities, lengths and number of foci as indicated in the figure legends. The χ^2^ test was used to compare the TUNEL and stage distributions as indicated in the figure legends. All tests and p- values (ns, not significant. *P < 0.05, **P < 0.01, ***P < 0.001, ****P < 0.0001) are provided in the corresponding legends and/or figures. Except for H3pS10 staining, which were performed on a single mouse per genotype, all other image analyses were performed on at least two independent mice for each genotype (see Table S2 – raw data).

## Data availability

All unique reagents generated in this study are available from the lead contact with a completed Materials Transfer Agreement.

All image analyses were performed using Fiji/ImageJ 1.53t, with the “MeiQuant” set of tools ^67^ available on github (https://github.com/MontpellierRessourcesImagerie/meiosis_bar).

Any additional information required to reanalyze the data reported in this paper is available from the lead contact upon request.

## Acknowledgement

We thank our laboratory members and Frederic Baudat for helpful discussion and reading. We thank Julie Chaumeil and David Lleres for their advices and protocol for immuno-FISH experiments. We thank Joao Matos for fruitful discussions and exchange of material. We thank Tatsuya Hirano for anti NCAPD2 used in immunofluorescence experiments. We thank Bernard de Massy and Christine Brun for the anti-REC114 antibody, Attila Tóth for the anti-HORMAD1 and -IHO1 antibodies, Alberto M Pendas for the anti-REC8 and -RAD21L antibodies and Bolcun-Filas’ laboratory for the anti-H1t antibody. We thank Ignasi Roig for anti-TRIP13 antibody reference. We thank Marie-Pierre Blanchard, MRI facility, for her help with STED microscopy. We thank the following Montpellier Biocampus facilities for their service: Réseau des Animaleries de Montpellier (RAM) for animal care, Réseau d’Histologie Expérimentale de Montpellier (RHEM) for histology and Montpellier Resources Imagerie (MRI) for microscopy.

The Centre for Structural Biology (CBS) is a member of France-BioImaging (FBI) and the French Infrastructure for Integrated Structural Biology, two national infrastructures supported by the French National Research Agency (grant n. ANR-10-INBS-04-01 and ANR-10-INBS-05, respectively). TR’s group is supported by the CNRS INSERM ATIP-Avenir 2017 program, ANR CONDENSin3R (ANR- 20-CE12-0016-02), ANR COMORE (ANR-24-CE12-2240-01), and La Ligue Contre le Cancer grants. LDT is funded by a PhD studentship from Université Montpellier, UM, ED CBS2, and a fourth year PhD studentship from Fondation pour la Recherche Médicale (FRM). AN is funded by a postdoctoral fellowship from Fondation pour la Recherche Médicale (FRM).

## Authors contribution

LDT performed most of the experiments, analyzed data and wrote the manuscript. BD performed experiments for *Ncapd2* cKO mutant establishment and initial analysis. AN performed experiment for FISH and IF analysis. AB performed MLH1 staining/counting, and staging. EG performed TUNEL analysis. JC developed ImageJ tools and participate to the image analysis. TR conceived the project, designed and performed experiments, analyzed data, and wrote the manuscript.

## Declaration of interests

The authors declare no competing interests.

## Extended data

Supplementary discussion.

Table S1: antibodies used for the study.

Figures S1 to S5.

## Notes

### Competing Interest Statement

The authors have declared no competing interest.

