## Extended Data for "Mammalian condensin I controls higher-order chromosome organization and homologous recombination in meiotic prophase I"

### Supplementary Discussion

***Ncapd2* cKO, a mouse genetic model to study condensin I in prophase I**. SMC-condensin complexes play a key role in organizing chromatin (Hirano, 2012). As in mammals the different condensin subunits are essential for embryonic development, we generated a *Ncapd2* cKO mouse line in which the gene encoding the NCAPD2 condensin I subunit is specifically ablated at the onset of prophase I. Using this model, we confirmed that NCAPD2 is essential for embryonic development and viability. This indicates that the formation of an active condensin I complex depends on the presence of this protein, and also that condensin I fulfils specific and essential functions during development. In meiotic prophase I, immunofluorescence analysis using an anti-NCAPD2 antibody suggests that the protein remains detectable in ~25% of early prophase I *Ncapd2* cKO nuclei, up to diplonema, where it became detectable in ~80% of nuclei. We think that incomplete or partial Cre-mediated excision leads to the presence of the protein in a fraction of mutant nuclei, which increase throughout prophase I. From this interpretation, we propose that NCAPD2 plays a pivotal role to proceed after pachynema. This explains the increase of mutant cells at this stage that we could interpret as an accumulation of cells hindered in their progression throughout prophase I. The late meiotic prophase I cells observed in the mutant correspond to pseudo-WT cells that express enough NCAPD2 to complete meiosis, thus explaining the presence of spermatozoa and subfertility. Nevertheless, we cannot exclude a role of condensin I in late prophase I, undetectable in a population contaminated by cells expressing a WT level of NCAPD2. Alternatively, the signal detected in the *Ncapd2* cKO mutant might correspond to a non-specific signal, more pronounced at diplonema, implying that the condensin I prophase I function is not essential for prophase I completion and subsequently for sperm formation. Thus, for clarity, we excluded diplonema and later meiotic cells from our analysis. Unlike NCAPD2 depletion at the onset of prophase I in males (this study), depletion of condensin I at metaphase entry in females has no impact on fertility (Houlard et al., 2015). This highlights the specific prophase I role of condensin I in mammalian spermatocytes. Of note, the STRA8 promoter used in this study to induce the CRE recombinase has a male specific expression, which then excludes the possibility to study female potential prophase I defects for the moment.

| Antibody | Source | Application | Dilution |
|---|---|---|---|
| Guinea pig anti-SYCP3 | B. de Massy (Grey *et al*., 2009) | IF | 1/1000 |
| Rabbit anti-SYCP3 | Abcam, Ab15093 | IF, WB | 1/400, 1/1000 |
| Mouse anti-SYCP3 | Abcam, Ab97672 | IF | 1/400 |
| Rabbit anti-REC8 | A. Pendas (Cristina Gutiérrez-Caballero *et al.* 2011) | IF, WB | 1/400, 1/1000 |
| Rabbit anti-RAD21L | A. Pendas (Cristina Gutiérrez-Caballero *et al.* 2011) | IF, WB | 1/50, 1/1000 |
| Guinea pig anti-HORMAD1 | A. Toth (Wojtasz *et al*. 2009) | IF, WB | 1/400, 1/1000 |
| Rabbit anti-TRIP13 | Proteintech, 19602-1-1P | IF, WB | 1/500, 1/1000 |
| Rabbit anti-SYCP1 | Abcam, Ab15090 | IF, WB | 1/400, 1/1000 |
| Rabbit anti-SIX60S1 | A. Pendas (Laura Gómez-H *et al*., 2016) | IF | 1/50 |
| Rabbit anti-TEX12 | Proteintech, 17068-1-1P | IF | 1/100, 1/1000 |
| Rabbit anti-CHD1 | Invitrogen, PA5-82795 | IF, WB | 1/150, 1/1000 |
| Rabbit anti-MEI4 | From B. de Massy (Kumar *et al*., 2010) | IF | 1/50 |
| Mouse anti-γH2AX | Millipore, 05-636 | IF | 1/10000 |
| Rabbit anti-DMC1 | Santa Cruz, H100 | IF, WB | 1/200 |
| Mouse anti-MLH1 | BD Pharmagen, 551091 | IF, WB | 1/50, 1/500 |
| Guinea pig anti-HEI10 | Home made (De muyt *et al.* 2025) | IF, WB | 1/200, 1/500 |
| Rabbit anti-HEIP1 | Home made (De muyt *et al.* 2025) | IF | 1/200 |
| Guinea pig anti-H1T | Gift from E. Bolcun-Filas Lab. Jackson Laboratory | IF | 1/500 |
| Goat anti-NCAPD2 | Home made (this study) | WB | 1/1000 |
| Rabbit anti-NCAPD2 | T. Hirano (Lee *et al*. 2011) | IF | 1/200 |
| Rabbit anti-β-Tubulin | Abcam, Ab6046 | WB | 1/5000 |
| Rabbit anti-H3pS10 | Sigma 06-570 | IF | 1/500 |
| Anti-Guinea Pig Alexa 488 | Jackson immunoresearch, 706-545-148 | IF | 1/250 |
| Anti-Rabbit Alexa 488 | Fisher Scientific, A21206 | IF | 1/500 |
| Anti-Mouse Alexa 488 | Jackson immunoresearch, 715-225-150 | IF | 1/50 |
| Anti-Guinea Pig Alexa 555 | Jackson immunoresearch, 706-165-148 | IF | 1/250 |
| Anti-Rabbit Alexa 555 | Invitrogen, A21428 | IF | 1/500 |
| Anti-Mouse Alexa 555 | Abcam, Ab150110 | IF | 1/500 |
| Anti-Guinea Pig Alexa 647 | Jackson immunoresearch, 706-175-148 | IF | 1/400 |
| Anti-Rabbit Alexa 647 | Abcam, Ab150063 | IF | 1/500 |
| Anti-Mouse Alexa 647 | Invitrogen, A31571 | IF | 1/500 |
| Anti-Mouse StarORANGE | Abberior GMBH, STORANGE-1001-500UG | IF | 1/100 |
| Anti-Rabbit StarRED | Abberior GMBH, STRED-1006-500UG | IF | 1/100 |
| Anti-Guinea Pig StarRED | Abberior GMBH, STRED-1002-500UG | IF | 1/100 |
| Anti-Guinea Pig HRP | Jackson immunoresearch, 706-035-148 | WB | 1/5000 |
| Anti-Mouse HRP | Jackson immunoresearch, 715-005-150 | WB | 1/3000 |
| Anti-Goat HRP | Santa Cruz, Sc2354 | WB | 1/5000 |

**Table S1 : List of antibodies used for western blot (WB) and immunofluorescence (IF) experiments.**

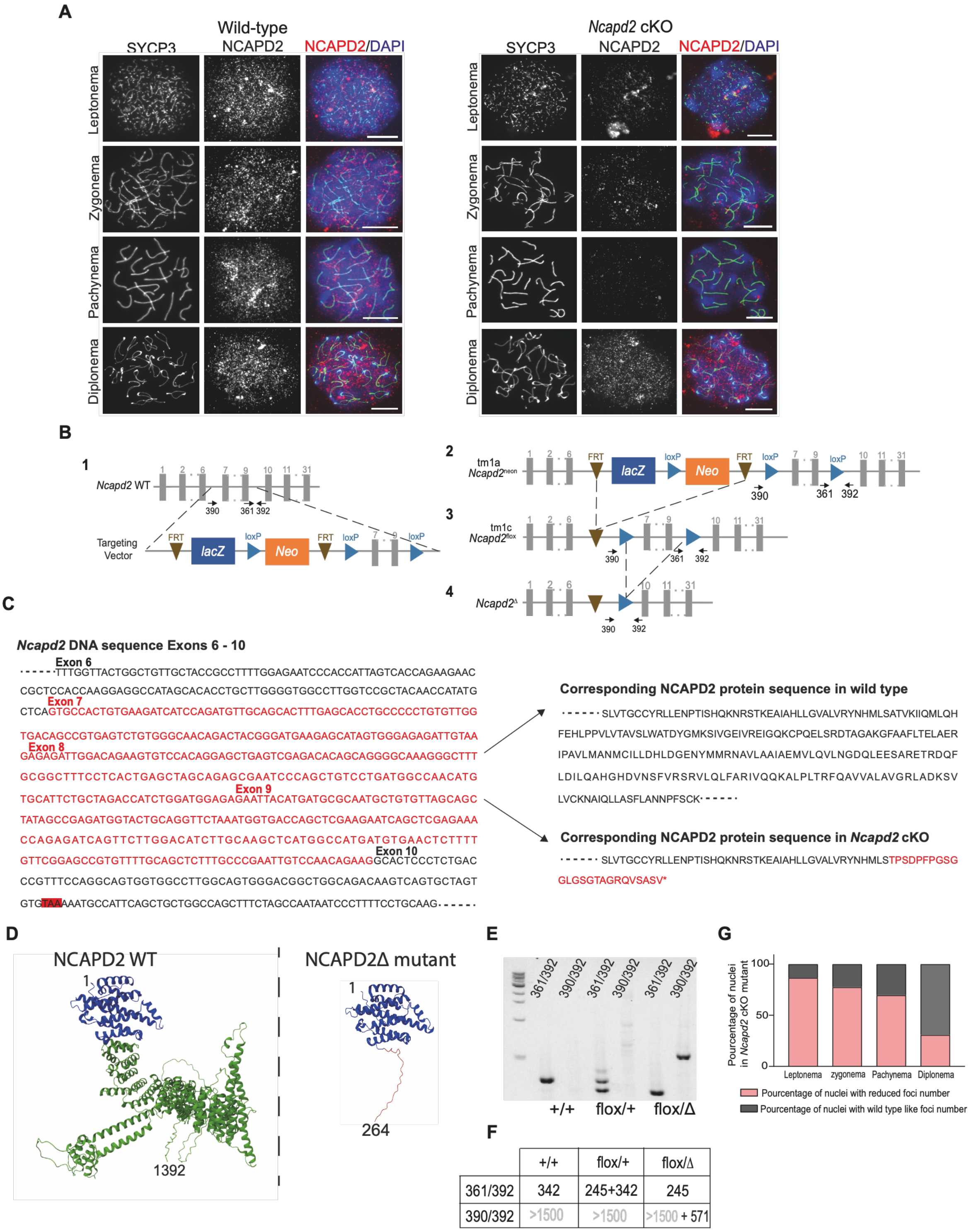

**Figure S1**

### Figure S1. Generation of *Ncapd2* cKO mice

(A) Representative images of SYCP3 (green) and NCAPD2 (red) immunostaining and DAPI staining (blue) in WT and *Ncapd2* cKO spermatocyte from leptonema to diplonema (18 dpp mice); scale bars, 10 μm.

(B) Schematic representation of the generation of *Ncapd2* cKO mice. (*1*) *Ncapd2* locus in WT ES cells with insertion in the intronic regions flanking exons 7 and 9 of the targeting vector. (*2*) Result of the insertion: generation of the *Ncapd2*$^{tm1a(EUCOMM)Wtsi\ (neon)}$ allele. (3) *Ncapd2*$^{flox}$ allele: locus after removal of the lacZ and neomycin cassettes. (4) *Ncapd2*$^{\Delta}$ allele: locus after loxP deletion by CRE. Grey boxes, exons; black arrows indicate the position of the genotyping primers. See Materials and Methods for more information.

(C) (*Left*) DNA sequence of the *Ncapd2* locus between exons 6 and 10 (exon start is annotated on the top of the sequence). In red: sequence removed by Cre recombinase-mediated excision. Highlighted in red: STOP codon that will lead to a premature translation arrest. (*Right*) Corresponding protein sequence in WT and *Ncapd2* cKO mice (NCAPD2Δ protein).

(D) AlphaFold 3 structural model of the NCAPD2 (1392 amino acids) and NCAPD2Δ (264 amino acids) proteins.

(E) Mouse genotyping. Primers 392 and 361 detect the WT and *Ncapd2*$^{flox}$ alleles. See Materials and Methods for primer sequences and panel (E) for expected PCR product sizes.

(F) Table showing the expected sizes of the PCR products using the different PCR primer pairs.

(G) Percentage of nuclei with reduced number or WT-like number of NCAPD2 foci in *Ncapd2* cKO spermatocytes. Nuclei were considered to have fewer NCAPD2 foci if their number of foci fell below the confidence interval of the WT distribution. Data are the mean of 2 independent experiments. Number of nuclei at leptonema: 26, 31 (wild type), 23, 22 (*Ncapd2* cKO); number of nuclei at early zygonema: 46, 48 (WT), 41, 36 (*Ncapd2* cKO); number of nuclei at pachynema: 68, 70 (WT), 40, 69 (*Ncapd2* cKO); number of nuclei at diplonema: 31, 22 (WT), 17, 16 (*Ncapd2* cKO).

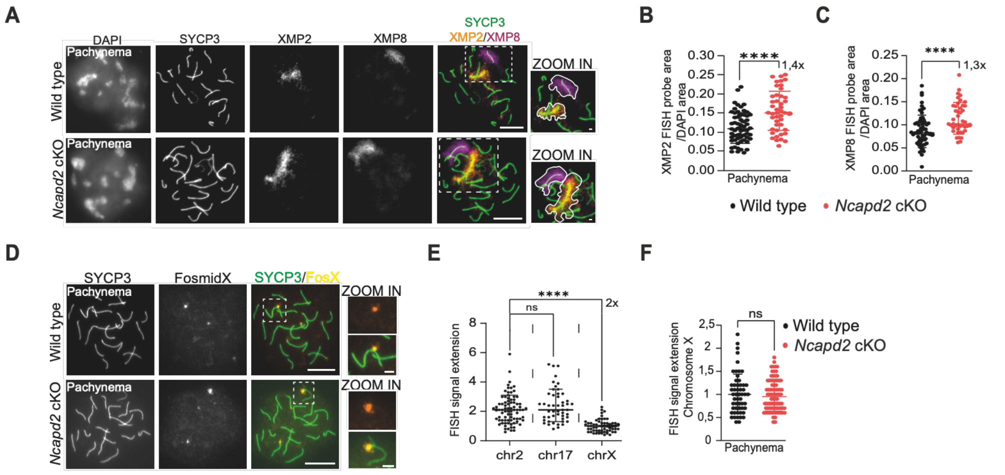

Figure S2

**Figure S2. NCAPD2 promotes chromatin compaction on chromosomes 2 and 8**

(A) Whole autosome chromatin staining. Representative images of WT and *Ncapd2* cKO pachytene nuclei immunostained with SYCP3 (green) and labelled with FISH probes for chromosomes 2 (XMP2, orange) and 8 (XMP8, magenta) in WT and *Ncapd2* cKO spermatocytes. The measured area is delimitated by a white border in the zoomed images.

(B and C) Quantification of the XMP2 (B) and XMP8 (C) FISH probe areas relative to the DAPI area in pachytene nuclei of WT and *Ncapd2* cKO spermatocytes. Number of pachytene nuclei: 62 (WT), 45 (*Ncapd2* cKO) for XMP2 and 63 (WT) and 43 (*Ncapd2* cKO) for XMP8.

(D) X chromosome labelled with fosmids. Representative images of WT and *Ncapd2* cKO pachytene nuclei immunostained with SYCP3 (green) and labelled with chromosome X fosmids (yellow). The length of the FISH signal extension is indicated with a white dashed line in the zoomed images.

(E) Quantification of the FISH signal extension on chromosomes 2, 17 and X (µm) in pachytene nuclei of WT mice. Number of pachytene nuclei: 81 (chromosome 2), 60 (chromosome 17), 59 (chromosome X).

(F) Quantification of the FISH signal extension on chromosome X (µm) in pachytene nuclei of WT and *Ncapd2* cKO spermatocytes. Number of pachytene nuclei: 59 (WT) and 82 (*Ncapd2* cKO).

For all experiments, 18 dpp mice were used. White boxes indicate regions highlighted in the zoomed images. Scale bars, 10 μm and 2 μm for zoomed images. P values were determined using the two-tailed unpaired Mann–Whitney test. Black or red bars show the mean values ± SD.

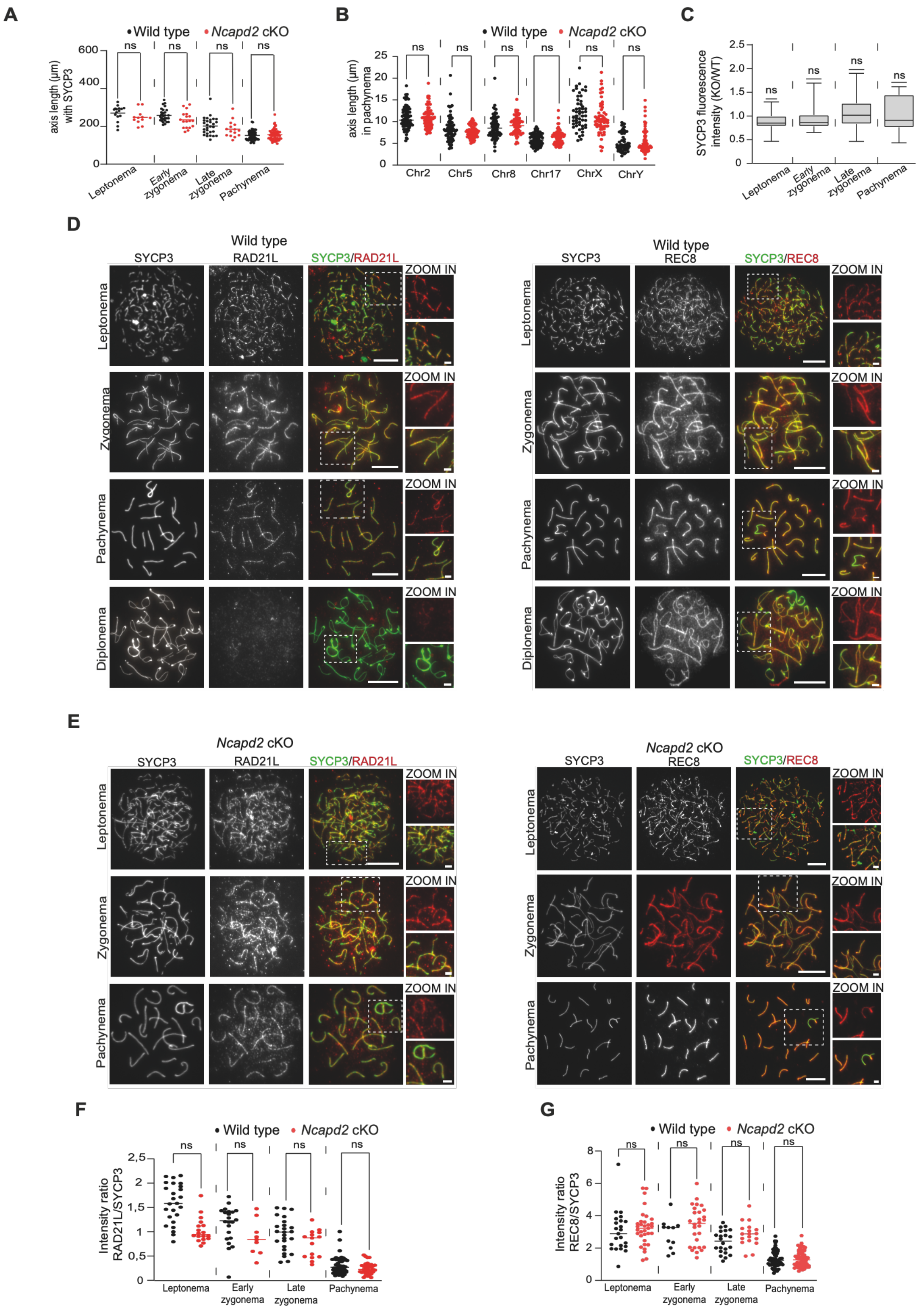
A
Wild type
Ncapd2 cKO
ns
axis length (µm) with SYCP3
600
400
200
0
Leptonema
Early zygonema
Late zygonema
Pachynema
B
axis length (µm) in pachynema
Chr2
Chr5
Chr8
Chr17
ChrX
ChrY
C
SYCP3 fluorescence intensity (KO/WT)
D
Wild type
SYCP3
RAD21L
SYCP3/RAD21L
ZOOM IN
Leptonema
Zygonema
Pachynema
Diplonema
REC8
SYCP3/REC8
E
Ncapd2 cKO
F
Intensity ratio RAD21L/SYCP3
G
Intensity ratio REC8/SYCP3

**Figure S3**

**Figure S3. SYCP3 and cohesin localization pattern are not affected in *Ncapd2* cKO spermatocytes**

(A) Quantification of the axis length (µm) using SYCP3 immunostaining from leptonema to pachynema in WT and *Ncapd2* cKO spermatocytes. Number of WT and *Ncapd2* cKO nuclei: leptonema: 16, 12; early zygonema: 30, 18; late zygonema: 29, 15; pachynema: 62, 58.

(B) Quantification of axis length (µm) using SYCP3 immunostaining on the chromosomes 2, 5, 8, 17, X and Y identified by FISH in pachytene nuclei of WT and *Ncapd2* cKO spermatocytes. Number of WT and *Ncapd2* cKO nuclei at pachynema: 83, 80 for chromosome 2; 81, 81 for chromosome 5; 83 and 80 for chromosome 8; 81 and 81 for chromosome 17; 56 and 54 for chromosome X; 59 and 54 for chromosome Y.

(C) Quantification of the SYCP3 signal intensity variation from leptotene to pachytene nuclei of *Ncapd2* cKO compared with WT. The ratios of the SYCP3 integrated intensity of *Ncapd2* cKO nuclei to the mean of the SYCP3 integrated intensity in WT nuclei at each stage are plotted. P values were determined using the one-sample t-test and a hypothetical value of 1. Number of nuclei at leptonema: 16; early zygonema: 18; late zygonema: 15; pachynema: 57.

(D) Representative images of (*left)* SYCP3 (green) and RAD21L (red) immunostaining and (*right*) SYCP3 (green) and REC8 (red) immunostaining from leptotene to diplotene nuclei of WT spermatocytes.

(E) Representative images of (*left*) SYCP3 (green) and RAD21L (red) immunostaining and (*right*) SYCP3 (green) and REC8 (red) immunostaining in *Ncapd2* CKO spermatocyte nuclei from leptotene to pachytene.

(F and G) Quantification of RAD21L (F) and REC8 (G) signal intensity relative to SYCP3 signal from leptonema to pachynema in WT and *Ncapd2* cKO mice. Number of WT and *Ncapd2* cKO nuclei at leptonema: RAD21L: 25, 18; REC8: 23, 33; early zygonema: RAD21L: 22, 9, REC8: 11 and 28; late zygonema: RAD21L: 25 and 15; REC8: 22, 18; pachynema: RAD21L: 54 and 36; REC8: 54 and 61.

All experiments were done with18 dpp mice. White boxes indicate regions highlighted in the zoomed images. Scale bars, 10 µm and 2 µm for zoomed images. (a, b, g and g): P values were determined using the two-tailed unpaired Mann–Whitney test. Black or red bars show the mean values ± SD.

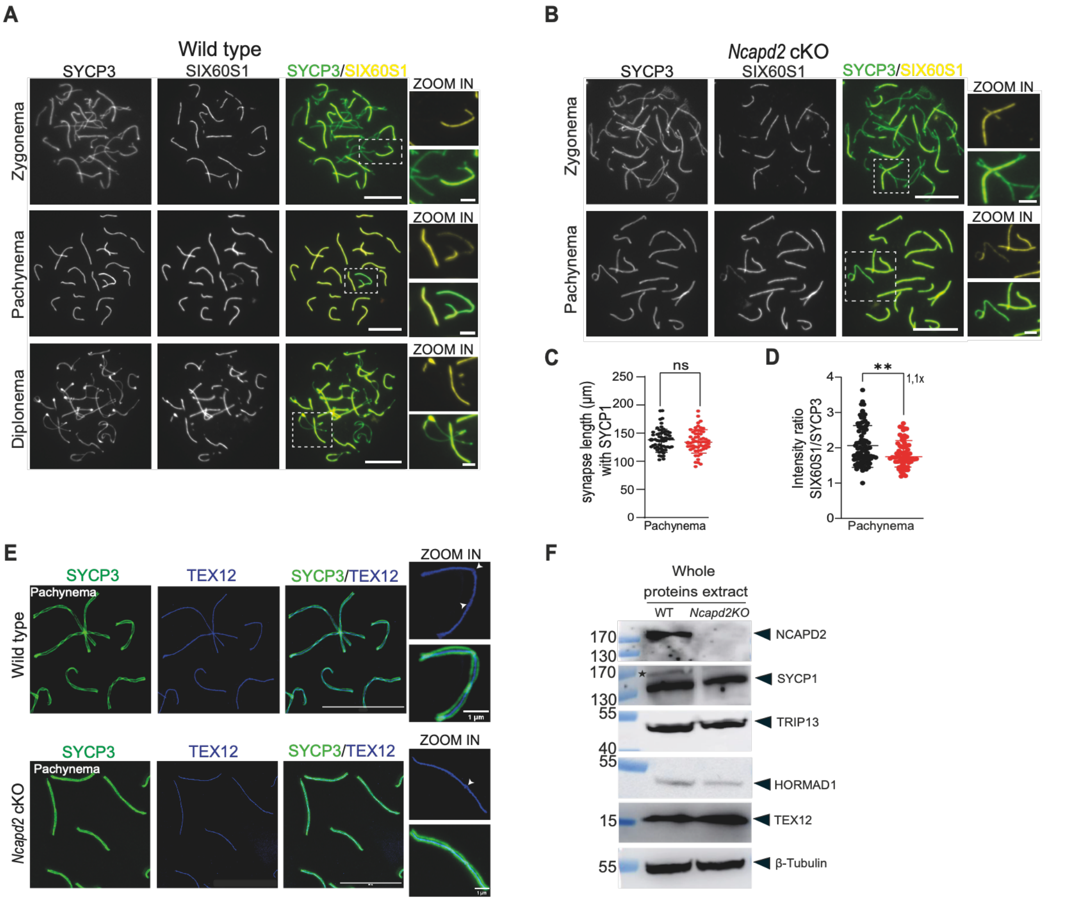

**Figure S4**

**Figure S4. NCAPD2 does not control synapsis length but controls SIX6OS1 loading.**

(A and B) Analysis of SIX6OS1 loading. Representative images of SYCP3 (green) and SIX6OS1 (yellow) immunostaining from zygotene to diplotene nuclei of WT (A) and in zygotene/pachytene nuclei of *Ncapd2* cKO spermatocytes (B).

(C) Measurement of synapsis length (µm) using SYCP1 immunostaining in pachytene nuclei of WT and *Ncapd2* cKO spermatocytes. Number of nuclei at pachynema: 61 (WT), 58 (*Ncapd2* cKO).

(D) Ratio of SIX60S1 signal intensity to SYCP3 in pachytene nuclei of WT and *Ncapd2* cKO spermatocytes. Number of nuclei at pachynema: 80 (WT) 61 (*Ncapd2* cKO).

(E) Representative STED microscopy images of SYCP3 (green) and TEX12 (blue) immunostaining in pachytene nuclei of WT and *Ncapd2* cKO mice.

(F) SC protein expression in *Ncapd2* cKO testes. Western blotting of whole protein extracts from testes of WT and *Ncapd2* cKO mice synchronized by retinoic acid performed with primary antibodies against: NCAPD2, SYCP1, TRIP13, HORMAD1, TEX12 and β-tubulin (loading control) (see Table S1). Expected molecular weights: NCAPD2 (~150 kDa), SYCP1 (~116 kDa), TRIP13 (~48 kDa), HORMAD1 (~45 kD1), TEX12 (~14 kDa) and β-tubulin (~55 kDa). n = 3 mice per genotype for NCAPD2, SYCP1 and HORMAD1 and n = 2 mice per genotype for TRIP13 and TEX12. The asterisk indicates a non-specific band.

Immunofluorescence experiments were performed using 18 dpp mice. White boxes indicate regions highlighted in the zoomed regions. Scale bars, 10 μm and 2 μm for zoomed images, but for (e) 1μm on zoomed images. P values were determined using the two-tailed unpaired Mann–Whitney test. Black or red bars show the mean values ± SD.

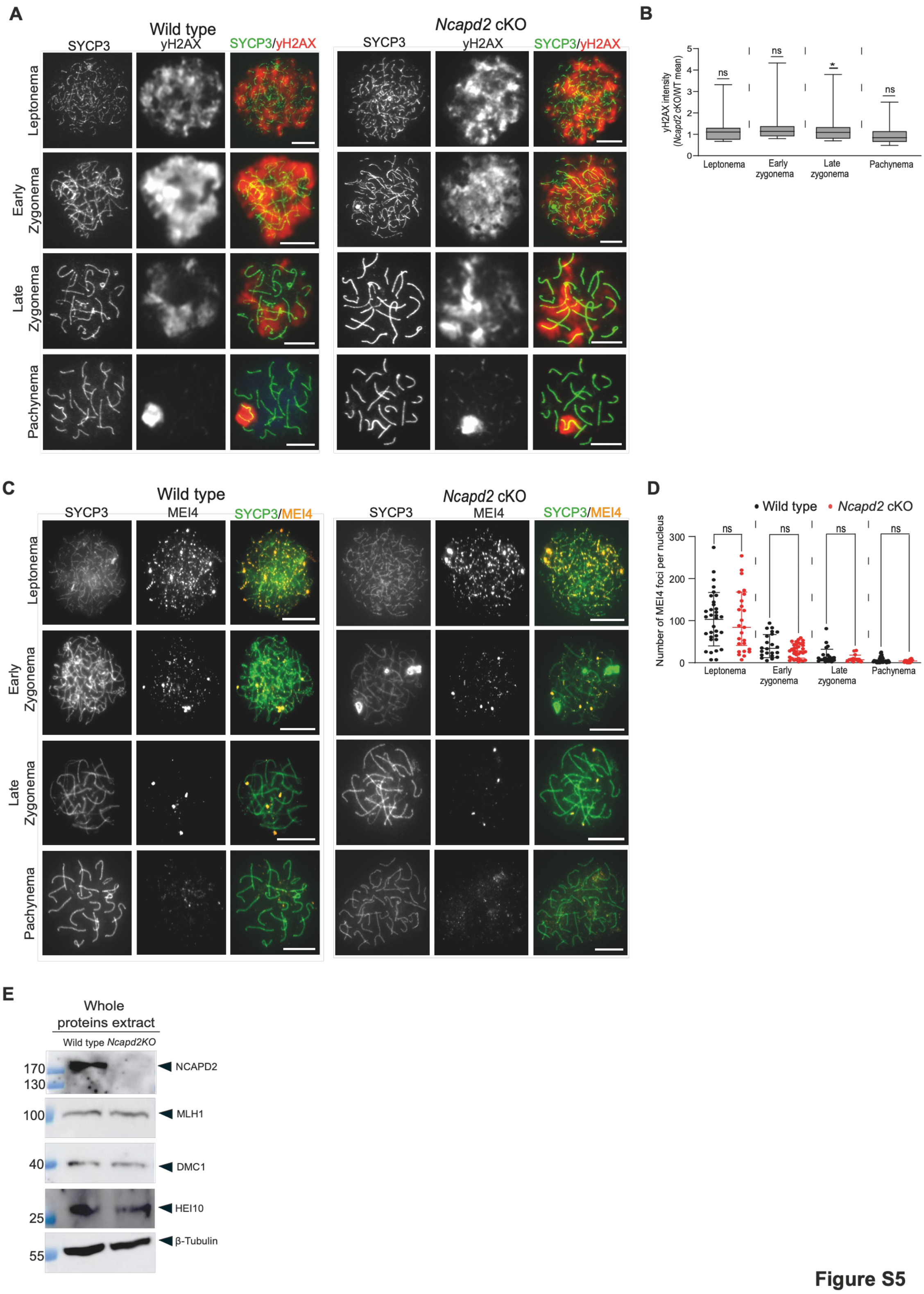
A
Wild type
Ncapd2 cKO
SYCP3
γH2AX
SYCP3/γH2AX
Leptonema
Early Zygonema
Late Zygonema
Pachynema
B
γH2AX intensity (Ncapd2 cKO/WT mean)
ns
ns
*
ns
Leptonema
Early zygonema
Late zygonema
Pachynema
C
MEI4
SYCP3/MEI4
D
• Wild type • Ncapd2 cKO
Number of MEI4 foci per nucleus
E
Whole proteins extract
Wild type Ncapd2KO
170
130
100
40
25
55
NCAPD2
MLH1
DMC1
HEI10
β-Tubulin

**Figure S5**

**Figure S5. NCAPD2 does not control DSB formation and MEI4 loading**

(A) Representative images of SYCP3 (green) and γH2AX (red) immunostaining of WT and *Ncapd2* cKO nuclei from leptotene to pachytene nuclei. Scale bar,10 μm.

(B) Quantification of γH2AX signal intensity in *Ncapd2* cKO spermatocyte nuclei. Graph shows the ratios of the γH2AX integrated intensity in *Ncapd2* cKO nuclei to the mean γH2AX integrated intensity in WT nuclei. P values were determined using the one-sample t-test and a hypothetical value of 1. Number of nuclei at leptonema: 18; early zygonema: 15; late zygonema: 29; pachynema: 86.

(C) Representative images of SYCP3 (green) and MEI4 (orange) immunostaining in WT and *Ncapd2* cKO spermatocyte nuclei from leptotene to pachytene Scale bar, 10 μm.

(D) Quantification of MEI4 foci from leptotene to pachytene nuclei of WT and *Ncapd2* cKO spermatocytes. Number of WT and *Ncapd2* cKO nuclei at leptonema: 33 and 25; early zygonema: 22 and 35; late zygonema: 24and 17; pachynema: 31 and 30. P values were determined using the two-tailed unpaired Mann–Whitney test.

(E) HR protein expression in *Ncapd2* cKO mice. Western blotting of whole protein extracts from testes of WT and *Ncapd2* cKO mice synchronized by retinoic acid using primary antibodies against NCAPD2, MLH1, DMC1, HEI10 and β-tubulin (loading control) (see Table S1). Expected molecular weights: NCAPD2 (~150 kDa), MLH1 (~85 kDa), DMC1 (~38 kDa), HEI10 (~31 kD1) and β-tubulin (~55 kDa); n = 3 mice per genotype for NCAPD2, MLH1 and DMC1 and n = 1 mouse per genotype for HEI10.

Immunofluorescence experiments were performed using 18 dpp mice. Scale bar, 10 μm. P values were determined using the two-tailed unpaired Mann–Whitney test. Black or red bars show the mean values ± SD.
